# SPEF1 mediates assembly of the central pair microtubule complexes in cilia of *Tetrahymena*

**DOI:** 10.64898/2025.12.03.689828

**Authors:** Mayukh Guha, Krishna Kumar Vasudevan, Yu-Yang Jiang, Panagiota Louka, Neeraj Sharma, Mireya Parra, Karl F. Lechtreck, Raphaël F.-X. Tomasi, Charles N. Baroud, Pascale Dupuis-Williams, Gerard W. Dougherty, Heymut Omran, Courtney Ozzello, Chad G. Pearson, Ewa Joachimiak, Dorota Wloga, Zachary W. Nurcombe, Corbin Black, Avrin Ghanaeian, Lina Mougharbel, Thomas M. Kitzler, Khanh Huy Bui, Jacek Gaertig

**Author notes:** correspondence (K.H.B), (J.G.).

## Abstract

SPEF1 is a conserved protein that reinforces the seam of central microtubules within the distal segment of motile cilia. We show that in *Tetrahymena*, loss of SPEF1 destabilized the central pair throughout the cilium length, producing gaps in the microtubule lattice, loss of proximal microtubule portions, and loss of lateral projections. While both central microtubules were affected, C1 suffered more damage than C2 and the defects were already apparent during ciliary assembly. *Tetrahymena* cells lacking SPEF1 had fewer cilia, which contained an excessively dense ciliary matrix and abnormal densities on the doublet microtubules. Although SPEF1 loss visibly affected only the central microtubules, it localized to all stable microtubules in the cell, both ciliary and non-ciliary. Within cilia, SPEF1 was enriched near the distal tip, but was also present throughout the middle segment, where it was more concentrated along the central than the outer doublet microtubules. In live cilia, SPEF1 particles were mostly stationary and turned over slowly, some diffused but none moved by IFT. We propose that SPEF1 acts as a stabilizer of central microtubules in both the middle and distal segment and its activity contributes to central microtubule assembly.

## Introduction

In most motile cilia, the axoneme contains nine outer doublet microtubules that surround a pair of singlet microtubules constituting the central pair (CP) apparatus (reviewed in (Loreng and Smith, 2017; Samsel et al., 2021)). In contrast, the non-motile (sensory) cilia typically lack the CP and the protein complexes that bend the axoneme, including radial spokes and dynein arms (reviewed in (Fisch and Dupuis-Williams, 2011)). Interactions between the CP and radial spokes control the activity of dynein arms that are attached to the outer doublet microtubules (Huang et al., 1982; Oda et al., 2014; Porter et al., 1992; Smith and Yang, 2004; Zhao et al., 2025). A loss of CP microtubules results in cilia paralysis (Dutcher et al., 1984; Mitchell and Sale, 1999; Rupp et al., 2001; Smith and Lefebvre, 1996; Smith and Lefebvre, 1997b; Warr et al., 1966). In humans, pathogenic variants in the CP protein components cause primary ciliary dyskinesia (Bautista-Harris et al., 2000; Olbrich et al., 2012; Stannard et al., 2004; Wohlgemuth et al., 2024). Importantly, the central and outer microtubules differ in the structure, mode of nucleation and protein composition (reviewed in (Loreng and Smith, 2017; Samsel et al., 2021)). Unlike the outer axonemal microtubules, the CP microtubules are not extensions of the basal body microtubules and their assembly requires katanin, and the Camsap3/Wdr47 complex (Dymek et al., 2004; Dymek and Smith, 2012; Lechtreck et al., 2013; Liu et al., 2021; Robinson et al., 2020; Saito et al., 2021; Sharma et al., 2007), which regulate the minus ends of non-centrosomal cytoplasmic microtubules (Buijs et al., 2021; Jiang et al., 2018; Jiang et al., 2014). As the CP and outer microtubules elongate, they become decorated with distinct sets of microtubule-associated proteins (MAPs) (Carbajal-González et al., 2013; Gui et al., 2022; Han et al., 2022; Kubo et al., 2023a; Li et al., 2022; Ma et al., 2019). Moreover, the two CP microtubules, C1 and C2, are decorated by different sets of MAPs (Carbajal-González et al., 2013; Gui et al., 2022; Han et al., 2022; Zhao et al., 2019; Zhu et al., 2025). Little is known about how MAPs are targeted to subtypes of axonemal microtubules. There is no indication that the central and outer microtubules are made of different isotypes or isoforms of tubulin (Gaertig and Wloga, 2008; Hoyle et al., 2008). On the other hand, within the same axoneme, the CP is more sensitive than the outer microtubules to certain tubulin mutations, including those in the flexible C-terminal tails (Kubo et al., 2023b; Nielsen et al., 2001; Thazhath et al., 2002). Overall, the mechanisms that differentiate the CP and outer microtubules remain obscure. SPEF1 is a conserved ciliary protein (Chan et al., 2005; Ostrowski et al., 2002) that selectively contributes to the CP microtubule assembly or stability. Loss-of-function alleles of SPEF1 in *Drosophila* (Wu et al., 2025) or knockdown of SPEF1 expression in mammalian ependymal cells (Zheng et al., 2019) result in axonemes that, on cross-sections, lack one or both central microtubules. Recombinant SPEF1 binds to microtubules *in vitro* and its overexpression *in vivo* bundles microtubules and stabilizes them against depolymerization (Dougherty et al., 2005; Gray et al., 2009; Legal et al., 2025; Werner et al., 2014; Zheng et al., 2019). In most ciliated cell types studied, SPEF1 is more concentrated within the ciliary distal segment (Chan et al., 2005; Konjikusic et al., 2023; Legal et al., 2025). Recently, cryo-electron tomography revealed that within the distal segment of cilia in *Tetrahymena*, SPEF1, through its CH domain, binds to both CP microtubules at the microtubule seam (Legal et al., 2025). Furthermore, SPEF1 dimers may cross-link the two central microtubules to control their alignment (Legal et al., 2025). SPEF1 is also present on some non-ciliary microtubules, including 15 protofilament microtubules in the pillar cells of the mammalian cochlea (Dougherty et al., 2005). In the *Xenopus* embryos, SPEF1 stabilizes microtubules at the leading edge of epithelial cells during radial intercalation (Werner et al., 2014). Furthermore, SPEF1 colocalizes with microtubules associated with the base of cilia, including the ciliary rootlets in the multiciliated cells of *Xenopus* (Werner et al., 2014) and the microtubule quartet bundle in *Trypanosoma brucei* (Gheiratmand et al., 2012; Pham et al., 2022). Here, we further investigate the significance of SPEF1 in *Tetrahymena,* a ciliated protist that assembles a wide array of diverse microtubules within a single cell. We find that in *Tetrahymena*, SPEF1 decorates most microtubule types, including both ciliary (central and outer) and non-ciliary microtubules. Although this broad distribution suggests that SPEF1 functions as a general microtubule-stabilizing protein, its loss profoundly affects only the CP, leading to the loss of portions of microtubules or projections. Although within the cilium, SPEF1 is highly enriched in the distal region, its absence dramatically disrupts CP organization throughout the middle segment, which constitutes most of the ciliary length. Live imaging further reveals that SPEF1 forms immobile particles within both the distal and middle segment, suggesting that, despite its distal enrichment, SPEF1 may promote CP stability along the entire ciliary shaft.

## Results

### *Tetrahymena* cells lacking SPEF1 assemble fewer cilia that exhibit abnormal motility

In *Tetrahymena thermophila,* there are two genes encoding paralogs of SPEF1 *TTHERM_00939230/SPEF1A* and *TTHERM_00161270*/*BBC31/SPEF1B.* Using mass spectrometry, SPEF1A was detected in cilia (Kubo et al., 2023a; Louka et al., 2018) and SPEF1B was detected in the basal bodies (Kilburn et al., 2007). In *Tetrahymena*, the CH domain of SPEF1 is bound along the seam of the two central microtubules within the ciliary tip segment (Legal et al., 2025). To determine the *in vivo* significance of SPEF1, we disrupted either *SPEF1A* or *SPEF1B* or both. The single gene knockout cells, SPEF1A-KO and SPEF1B-KO, swam 35% and 25% more slowly than the wild type, respectively. The double knockout cells (SPEF1AB-KO), were 56% slower than the wild type (Fig. 1C-E). Deletion of both SPEF1 paralogs also reduced the rate of phagocytosis, which requires the motility of oral cilia to filter food particles (Fig. 1H-K) and slowed culture growth (Fig. S1A). Unexpectedly, cells that lacked one or both SPEF1 paralogs assembled 30% fewer cilia as compared to the wild type (Fig. 1A,B and S1B). In the SPEF1AB-KO cells, the spacing between adjacent basal bodies within the ciliary rows appeared normal (Fig. 1A,B); therefore, the loss of SPEF1 likely increases the fraction of unciliated basal bodies. The length of cilia was unaffected (Fig. S1C). After deciliation, the SPEF1AB-KO cells regrew cilia with the elongation rate similar to the wild type (Fig. S1D), further indicating that the reduction in the number of cilia per cell is a result of fewer basal bodies undergoing ciliation.

**Figure 1.**
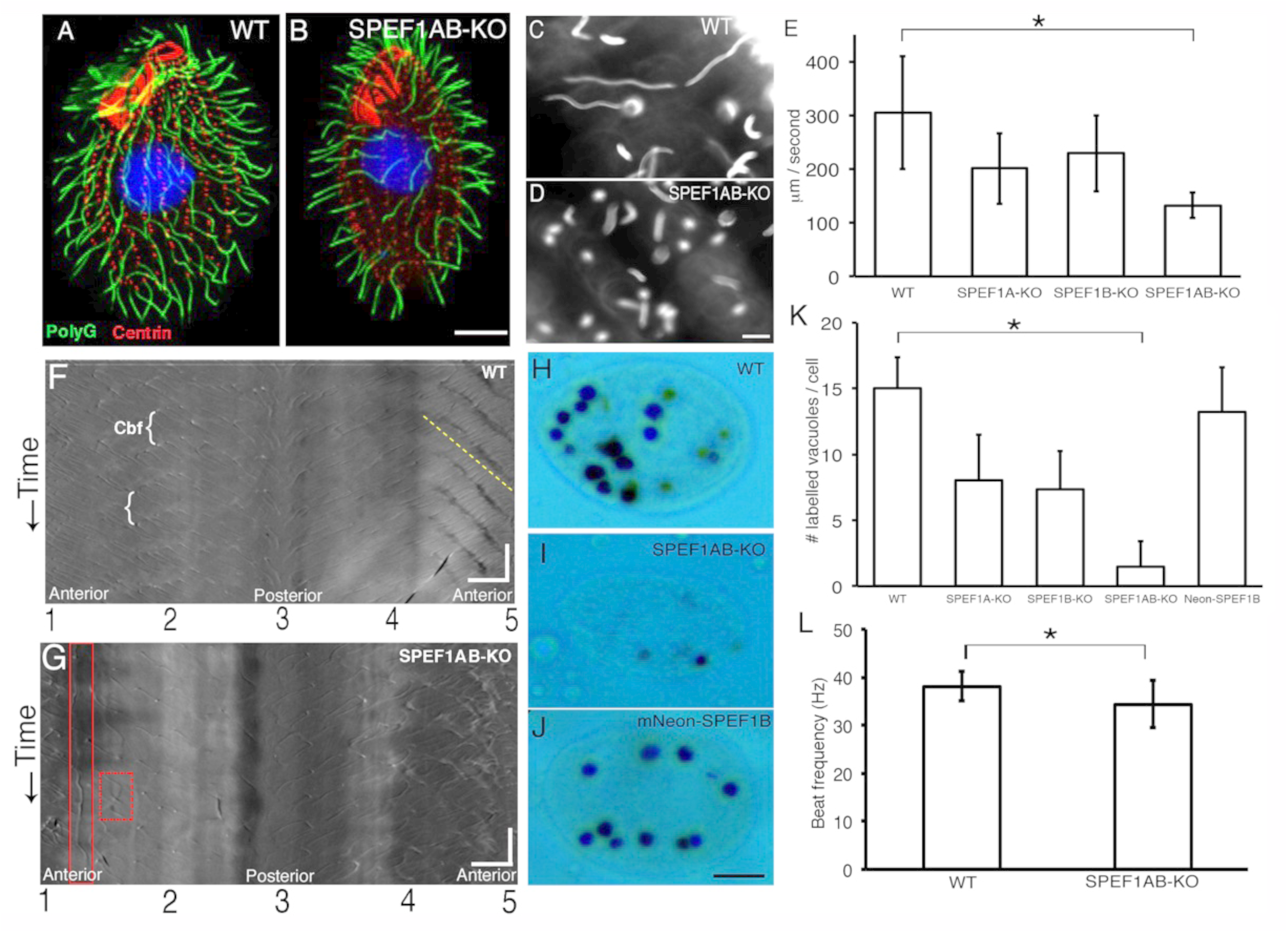
*Tetrahymena* cells lacking SPEF1 have fewer and abnormal cilia. (A,B) Confocal immunofluoroscence images of wild-type (A) and SPEF1AB-KO (B) cells stained with antibodies against polyglycylated tubulin (green) and centrin (red). Nuclei were stained with DAPI (blue). Scale bar = 10 μm. (C-D) Dark-field images of paths of swimming cells (1 sec exposure). Scale bar = 0.1 mm. (E) Cell swimming speeds. Mean ± SD μm/sec. WT: 305 ± 105 (N = 35); SPEF1A-KO: 201 ± 66 (N = 184); SPEF1B-KO: 229 ± 71 (N = 201); SPEF1AB-KO: 132 ± 24 (N = 51). *P < 0.0001. (F-G) Chronographs obtained from videos of free-swimming *Tetrahymena* cells (see Figure S2 for method details). The images represent an optical slice around the surface of a wild-type (F) and a SPEF1AB-KO (G) cell. The numbers (1 to 5) on the x-axis mark the positions around the cell circumference, with the split at the anterior cell tip between positions 1 and 5. The y-axis represents the passing time from top to bottom. The white brackets in the wild type (F) are an example of a single cilia beat frequency (cbf) measurement. Metachronal waves are visible as darker diagonal lines on the right anterior side of the wild-type cell surface (F) that are perpendicular to the diagonal lines representing individual cilia (dashed yellow line). In the SPEF1AB-KO chronogram (G), note a stalled cilium (red solid box) and a cilium beating with the power stroke in the opposite direction (red dashed box). Scales bars: x-axis = 10 μm, y-axis = 50 msec. (H-J) Bright field images of wild-type (H), SPEF1AB-KO (I) and mNeonGreen-SPEF1B rescue of SPEF1AB-KO (J) cells fed with India ink for 30 min. Scale bar = 10 μm. (K) Rates of phagocytosis quantified as an average number of ink-filled food vacuoles per cell in 30 min. Mean ± SD. WT: 15 ± 2.4 (N = 59); SPEF1A-KO: 8.1 ± 3.4 (N = 54); SPEF1B-KO: 7.4 ± 2.9 (N = 58); SPEF1AB-KO: 1.5 ± 2 (N = 88); mNeonGreen-SPEF1B rescue of SPEF1AB-KO: 13.2 ± 3.4 (N = 69). *P < 0.0001 (Tukey’s multiple comparisons test). (L) Ciliary beat frequencies (based on chronographs shown in F and G). Mean ± SD Hz. WT: 38.1 ± 3.2 (N = 45 cilia, N = 9 cells) and SPE1AB-KO: 34.4 ± 4.9 (N = 34 cilia, N = 6 cells). *P < 0.001 (two tailed t test). Abbreviations: cbf, cilium beat frequency.

Next, we performed high speed video recording of live wild-type and SPEF1AB-KO cells confined inside microfluidic channels (Movies 1 and 2). Space-time diagrams (Funfak et al., 2015) were used to examine the pattern and beat frequency of cilia across the entire cells (Fig. 1F,G; the short diagonal lines represent the traces of individual cilia, the white brackets connect subsequent traces of the same cilium over time and the distance between subsequent traces is used to calculate the beat frequency, see Fig. S2). The SPEF1AB-KO cilia showed a 10% reduction in the beat frequency (Fig. 1F,G,L) and displayed immotile cilia (Fig. 1G solid red box, yellow arrow in Movie 2) or cilia generating power strokes in the opposite direction (Fig. 1G, red dashed box). Near the anterior region of the wild-type cells, the metachronal waves are detectable as darker diagonal lines that are perpendicular to the diagonal short lines representing individual cilia (Fig. 1F, dashed yellow line (see (Funfak et al., 2015)). Such metachronal waves were not apparent in the SPEF1AB-KO cells (Fig. 1G). Overall, the diagonal line pattern of individual SPEF1AB-KO cilia was more variable, especially near the anterior cell end, indicating widespread but more subtle waveform defects (Fig. 1F,G). Thus, slow swimming exhibited by the SPEF1AB-KO cells is likely caused by both the reduced number and abnormal beating of cilia.

### *Tetrahymena* cells lacking SPEF1 have defective central pair microtubules in the middle ciliary segment

In the wild type *Tetrahymena*, transmission electron microscopy (TEM) revealed exclusively 9+2 cross-sections of the middle segment of axonemes (Fig. 2A,H). In SPEF1AB-KO, there were cross-sections lacking one (9+1, 22%) or both (9+0, 12%) central microtubules (Fig. 2B-D,I and L). The outer microtubules were unaffected. The cross-sections within the distal segment (recognized by the presence of one or more outer singlets) contained two central microtubules (Fig. 2J,K, wild type N = 9; SPEF1AB-KO N = 21), in agreement with the recent cryo-electron tomography (cryo-ET) study revealing only a subtle effect of SPEF1 loss on the CP microtubule alignment in the distal segment (Legal et al., 2025). Longitudinal sections of cilia revealed truncations of one of the two central microtubules at the proximal axoneme end, close to the basal body, and in the few cases analyzed, the truncated microtubule was located on the convex side of the ciliary bend (compare Fig. 2F,G with 2E). The two microtubules of the CP, designated C1 and C2, have projections of distinct size, periodicity and composition (Carbajal-González et al., 2013; Gui et al., 2022; Han et al., 2022; Linck et al., 1981; Zhu et al., 2025). C1 is typically positioned closer to the convex, while C2 is closer to the concave of the ciliary bend, respectively (Bernstein et al., 1994; Han et al., 2022; Lechtreck and Witman, 2007; Mitchell, 2003). In *Tetrahymena*, C1 is slightly longer and its minus end is embedded in an electron dense structure located immediately distal the transition zone, the axosome (Fig. 2E, white arrow marks the minus end of C1, white arrowhead marks the axosome). C2 is shorter and its minus end is not attached to the axosome (Fig. 2E) (Hausmann and Fischer-Defoy, 1978). The truncated microtubule in the SPEF1AB-KO cilia was positioned on the outside of the bend suggesting that it was C1 (Fig. 2F,G, white arrows). Unexpectedly, the non-truncated partner microtubule that was located on the inside of the bend, was connected to the axosome through a kink (Fig. 2F,G, empty black arrowheads). It appears therefore that the truncated microtubule is C1. It further appears that when C1 is missing, C2 connects to the axosome, taking the position that is normally occupied by C1. Because CP is critically important for ciliary motility (reviewed in (Loreng and Smith, 2017; Samsel et al., 2021)), the abnormal waveforms of SPEF1AB-KO cilia can be explained by the CP defects. However, we have also noticed that *Tetrahymena* mutants lacking CP protein components, tend to have fewer cilia. Namely, knockouts of genes encoding conserved CP MAPs: *PF16/SPAG6* and *PF20/SPAG16* (Adams et al., 1981; Smith and Lefebvre, 1996; Smith and Lefebvre, 1997a), both resulted in fewer cilia, except that unlike the SPEF1 knockout cilia, the PF16-KO and PF20-KO cilia were also shorter (Fig. S3). Cilia are also shorter in the *Tetrahymena* mutants lacking components of the C1b projection, SPEF2 and FAP69 (Joachimiak et al., 2021). Thus, the reduced ciliogenesis observed in the SPEF1AB-KO cells could be secondary to the CP defects.

**Figure 2.**
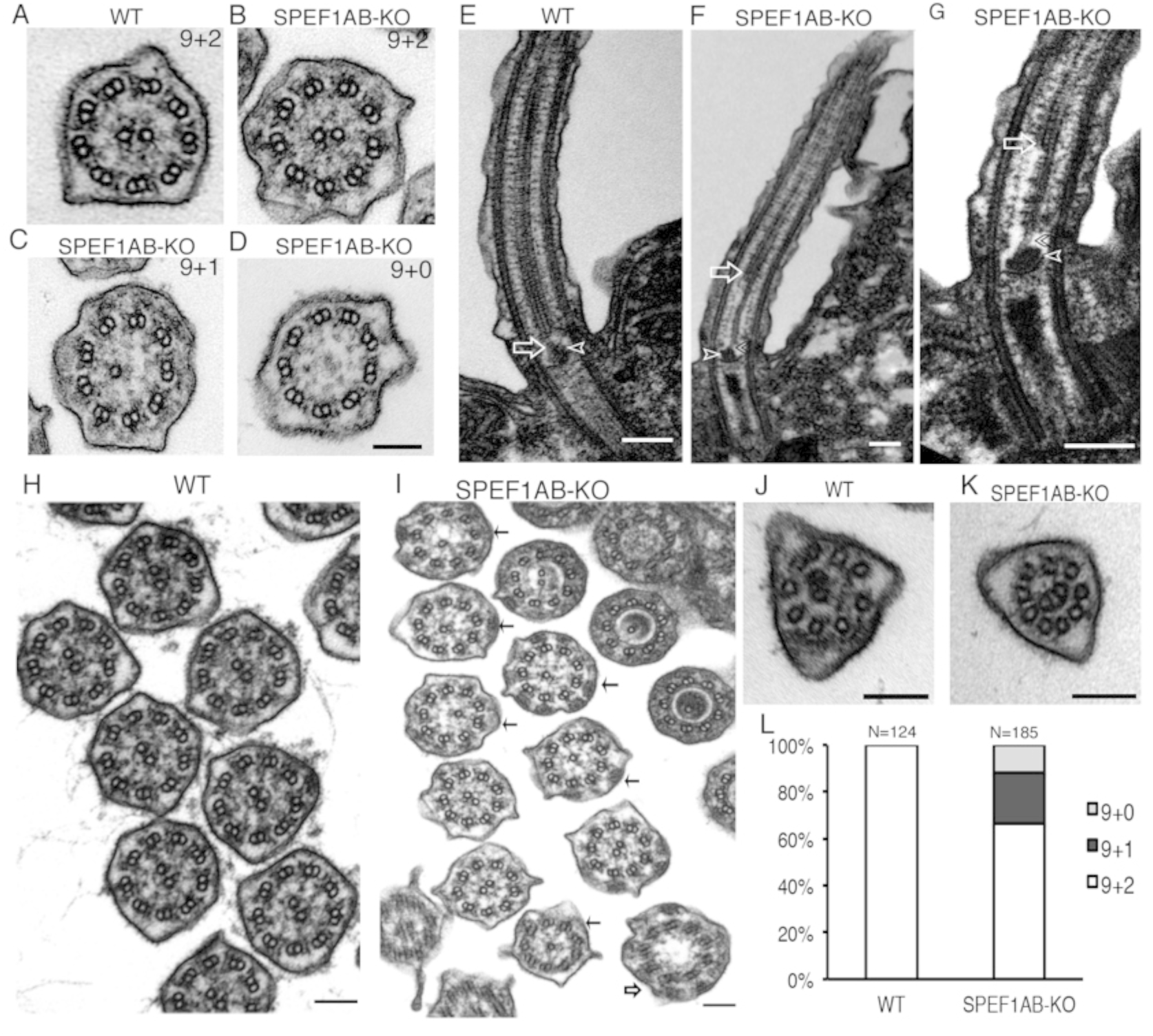
Central microtubule defects in the middle segment of cilia of *Tetrahymena* cells lacking SPEF1. (A-D) TEM cross-sections of the middle segment (doublet zone) of wild-type (A) and SPEF1AB-KO (B-D) cilia. Quantification of the CP microtubule defects is shown in panel L. Scale bar = 75 nm. (E-G) Longitudinal sections of a wild-type (E) and SPEF1AB-KO (F-G) locomotory cilia across the axoneme proximal region and basal body. Empty white arrows indicate the proximal end of C1 (in the mutant cilia identified based on its position on the outside of ciliary bend). Empty white arrowheads mark the axosome. Empty black arrowheads mark the kinked C2 end in SPEF1AB-KO. Scale bars = 150 nm. (H-I) TEM cross-section through groups of oral cilia (M rows). Abnormal middle segment configurations in SPEF1AB-KO (I) are marked (black arrows - 9+1, empty black arrows - 9+0). Scale bars = 100 nm. (J-K) Cross-sections through the distal ciliary segment (outer singlet zone). No obvious ultrastructural defects are seen in of SPEF1AB-KO (K). Scale bar = 100 nm. (L) Quantification of the doublet zone configurations in the middle segment cross-sections.

Given that SPEF1 localizes to both ciliary and non-ciliary microtubules in *Tetrahymena* and other ciliated species (see next section), we evaluated the effect of SPEF1 loss on the microtubules that form bundles within the cell cortex: longitudinal microtubule bundles (LMs) (positioned along the somatic ciliary rows) and transverse (TM) and postciliary (PC) microtubule rootlets, which are anchored at the basal bodies (reviewed in (Wloga and Frankel, 2012)). Judging by the average number of microtubule profiles, none of these cortical microtubule bundle types were affected in the SPEF1AB-KO cells (Table S2). Thus, despite its ubiquitous presence across the cell (see below), SPEF1 loss grossly affects only the CP microtubules.

SPEF1 is important for the CP organization in mammalian cultured cells (Zheng et al., 2019), *Drosophila* (Wu et al., 2025) and *Tetrahymena* (this study). To further examine the evolutionary conservation of SPEF1 function in motile cilia, we performed knockdown of the SPEF1 ortholog gene (*spef1*) using antisense morpholino oligonucleotides in *Danio rerio* (zebrafish) embryos. The *spef1* morphants exhibited phenotypes characteristic of ciliary dysfunction, including hydrocephalus, abnormal otolith development and abnormal L/R heart positioning (Fig. S4). These findings indicate that the ciliary role of SPEF1 is highly conserved, consistent with its fundamental role in CP organization.

### SPEF1 localizes to ciliary and non-ciliary cortical microtubules

To image SPEF1, we constructed a *Tetrahymena* strain with a GFP-SPEF1B transgene expressed in the SPEF1AB-KO background as follows. *Tetrahymena* strains lacking microtubule-associated proteins can have altered sensitivity to tubulin-binding drugs (Vasudevan et al., 2015; Wloga et al., 2009). Indeed, while the single knockouts and wild-type cells grew, the SPEF1AB-KO cells failed to multiply in the presence of 10 μM paclitaxel. We introduced transgenes expressing ectopic SPEF1B (mNeonGreen-SPEF1B or GFP-SPEF1B) into SPEF1AB-KO cells by selecting rescue transformants with 10 μM paclitaxel. Not only the paclitaxel hypersensitivity but also cilia-dependent phagocytosis (Fig. 1H-K) and culture growth defects (Fig. S1A) were rescued. Thus, SPEF1A and SPEF1B are partially functionally-redundant and the transgenic SPEF1B is sufficient to fulfill the function of total SPEF1. Using super-resolution structural illumination microscopy (SR-SIM) with the anti-acetyl-K40 tubulin antibody that labels cortical microtubules and cilia, and also total internal reflection fluorescence microscopy (TIRFM), we detected abundant mNeonGreen-SPEF1B in the cell cortex (Fig. 3A and Movies 3 and 4). mNeonGreen-SPEF1B highlighted the basal bodies and the basal body-associated microtubule bundles: TM and PC bundles (Fig. 3, yellow and white arrows, respectively). In the TM and PC bundles, the plus ends of microtubules are positioned away from the basal bodies (Thazhath et al., 2004). Interestingly, unlike in cilia, where SPEF1 is enriched near the microtubule plus ends at the distal tip (see below and (Legal et al., 2025)), no such pattern was seen in the basal body-associated microtubule bundles; to the contrary, mNeonGreen-SPEF1B seemed more abundant near the proximal (minus) ends (Fig. 3A insets on the right). The contractile vacuole pores (CVPs), containing bundles of unusual circular (pore) microtubules and associated rootlets, were strongly decorated with mNeonGreen-SPEF1B (Fig. 3A, blue arrow and Movie 3). In addition, TIRFM revealed a weak mNeonGreen-SPEF1B signal along the LM bundles that run parallel to the rows of somatic cilia (Movies 3 and 4, yellow arrows). Within cilia, mNeonGreen-SPEF1B was enriched near the distal tip (Fig. 3A, insets on the left), as shown earlier for SPEF1A-GFP expressed under the native promoter (see below and (Legal et al., 2025)), in agreement with the observations in *Xenopus* embryos (Gray et al., 2009; Konjikusic et al., 2023; Werner et al., 2014). However, in the multiciliated murine ependymal cells, SPEF1 is distributed uniformly along the cilium length (Wu et al., 2025; Zheng et al., 2019). We found that SPEF1 is enriched at the tips of cilia in human and murine multiciliated respiratory cells (Fig S5), indicating that the distal tip enrichment is conserved in some mammalian cell types.

**Figure 3.**
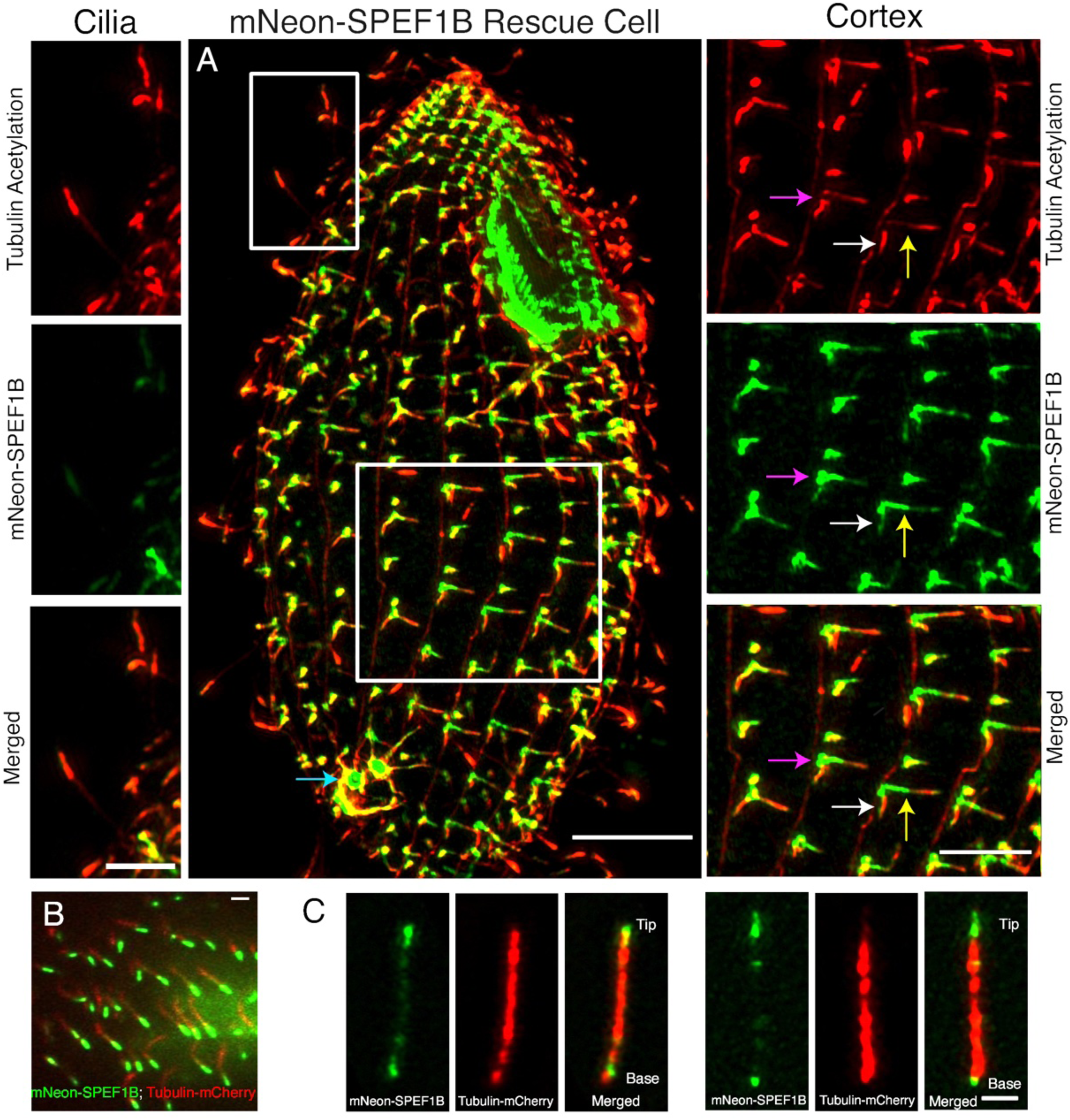
SPEF1 localizes to ciliary and non-ciliary (cortical) microtubules. **(A)** SR-SIM image of the SPEF1AB-KO cell phenotypically rescued by expression of mNeonGreen-SPEF1B (green), stained with an antibody against acetyl-K40 α-tubulin (red). mNeonGreen-SPEF1B is expressed under *MTT1* promoter without induction with cadmium ions. Boxes mark areas showing single channel images at higher magnifications, cilia (left) and the cell cortex (right). In the cell cortex, SPEF1 localizes to the basal bodies (magenta arrows), post-ciliary microtubules (white arrows), transverse microtubules (yellow arrows), contractile vacuole pores (blue arrow). Scale bars = 2.5 μm for insets and 5 μm for the whole cell panel. (B) A TIRFM image of a surface of a cell expressing mNeonGreen-SPEF1B (green) and β-tubulin-mCherry (red). (C) SR-SIM images of single cilia isolated from cells expressing mNeonGreen-SPEF1B (green) and β-tubulin-mCherry (red).

Next, we co-imaged mNeonGreen-SPEF1B with tagged β-tubulin (Btu2-mCherry) which confirmed that mNeonGreen-SPEF1B was highly enriched near the tips of cilia (Fig. 3B and Movie 5). SR-SIM of isolated cilia confirmed SPEF1B enrichment at the distal tip, but also revealed patches at the opposite end of the axoneme and also smaller foci scattered along the entire axoneme length (Fig. 3C). To see how the distribution of SPEF1 forms during cilia assembly, we imaged SPEF1A-GFP (expressed under native promoter in a wild type background) during vegetative growth with immunofluorescence using TAP952, a monoclonal antibody specific to monoglycylated (monoG) tubulin (Levilliers et al., 1995). In the mature full-length cilia, TAP952 labels strongly the distal tip and weakly the rest of the axoneme. In assembling cilia, TAP952 labels the entire cilium more uniformly (Brown et al., 1999). The full-length cilia invariably had a SPEF1A enriched distal segment (Fig. 4A-A’). Assembling cilia displayed a variable pattern, either already had a distally-enriched SPEF1A (Fig. 4A,A’, white arrows) or had SPEF1A scattered along the length or lacked SPEF1 foci (Fig. 4A,A’, yellow arrows). A similar pattern was observed in cells expressing GFP-SPEF1B (in the double KO background) regenerating cilia after pH shock deciliation (Fig. 4B-B’). Both patterns were seen in the nascent cilia of ∼ 1 μm in length. It therefore appears that the distal enrichment of SPEF1 forms early during cilia assembly, likely with a short delay from the time of initiation of cilium outgrowth from the basal body.

**Figure 4.**
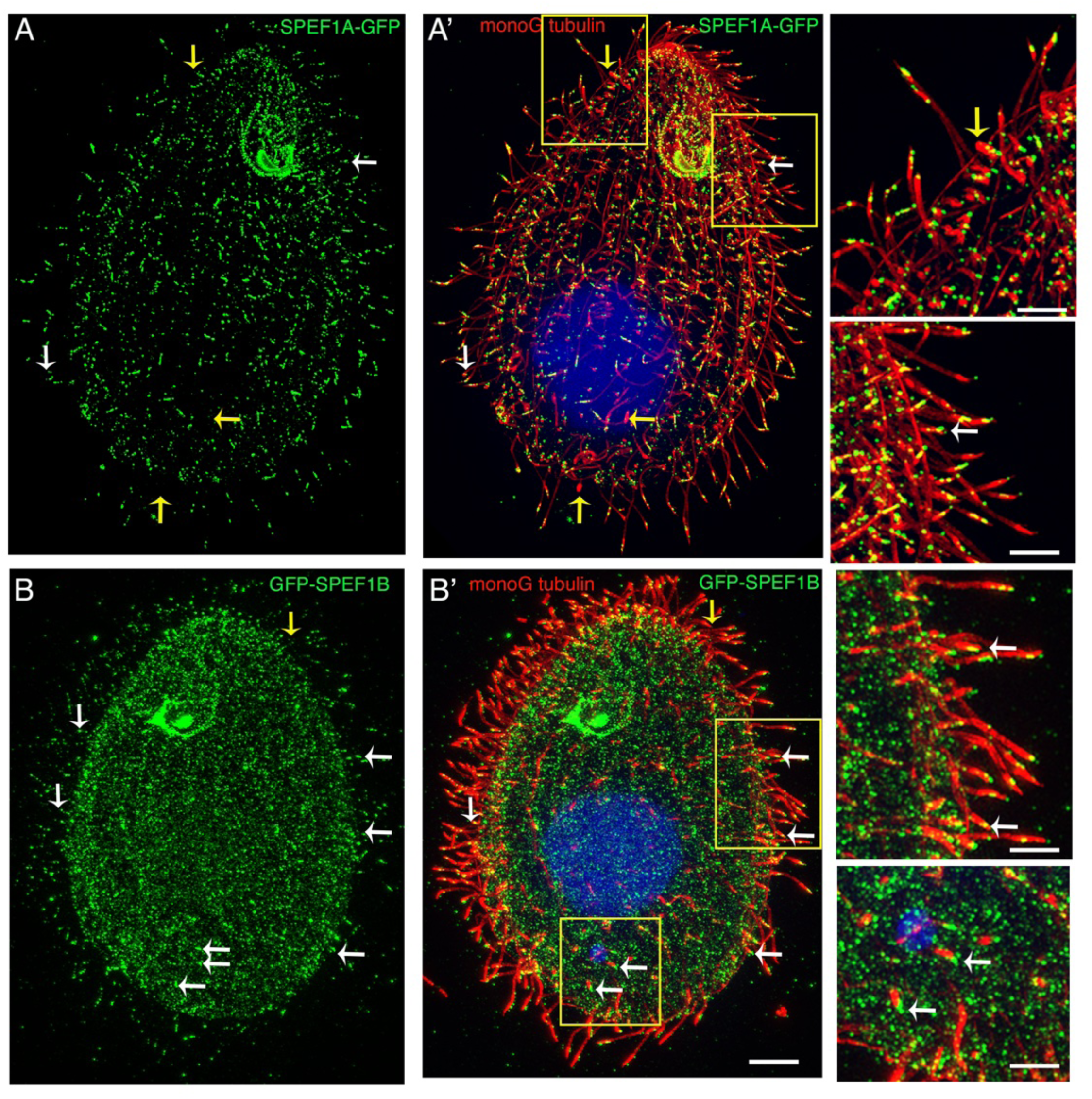
The ciliary distal tip enrichment of SPEF1 forms early during ciliary assembly. (A-B). SR-SIM images of cells expressing either SPEF1A-GFP under native promoter in the wild-type background (A) or GFP-SPEF1B under *MTT1* promoter (non-induced) in the SPEF1AB-KO background (B) (green) and labeled with TAP952 anti-monoglycylated tubulin antibody (red). The cell shown in (A) is from a standard vegetatively growing culture while the cell shown in (B) regenerates cilia 2 hr after pH shock deciliation. Boxes mark areas shown at a higher magnification in insets. White arrows point to the tips of short assembling cilia that already have a distally enriched SPEF1. Yellow arrows point to the tips of assembling cilia that either lack or have dispersed SPEF1 particles. Scale bars: 5 μm in the main panels and 2.5 μm in the insets.

### Within the middle segment, SPEF1 localizes to both the central and doublet microtubules

Given the selective effect of the loss of SPEF1 on the CP microtubules, we asked whether SPEF1 is targeted selectively to the CP microtubules. We tagged the C-terminus of natively-expressed SPEF1B with mCherry and studied its distribution by post-embedding immunoelectron microscopy (IEM) using anti-mCherry antibodies. In the middle ciliary segment, gold particles were present near both the central microtubules and outer doublet microtubules (Fig. 5A-D). Similar observations were made in negatively-stained ATP-reactivated axonemes (Fig. 5E,F) and quantification showed that gold particles were more frequently detected along the CP than along the doublets (Fig. 5G). Thus, SPEF1 is not exclusive to the distal tip compartment and within the middle segment, there is preference but not an absolute selectivity for association with the CP microtubules over the outer doublets.

**Figure 5.**
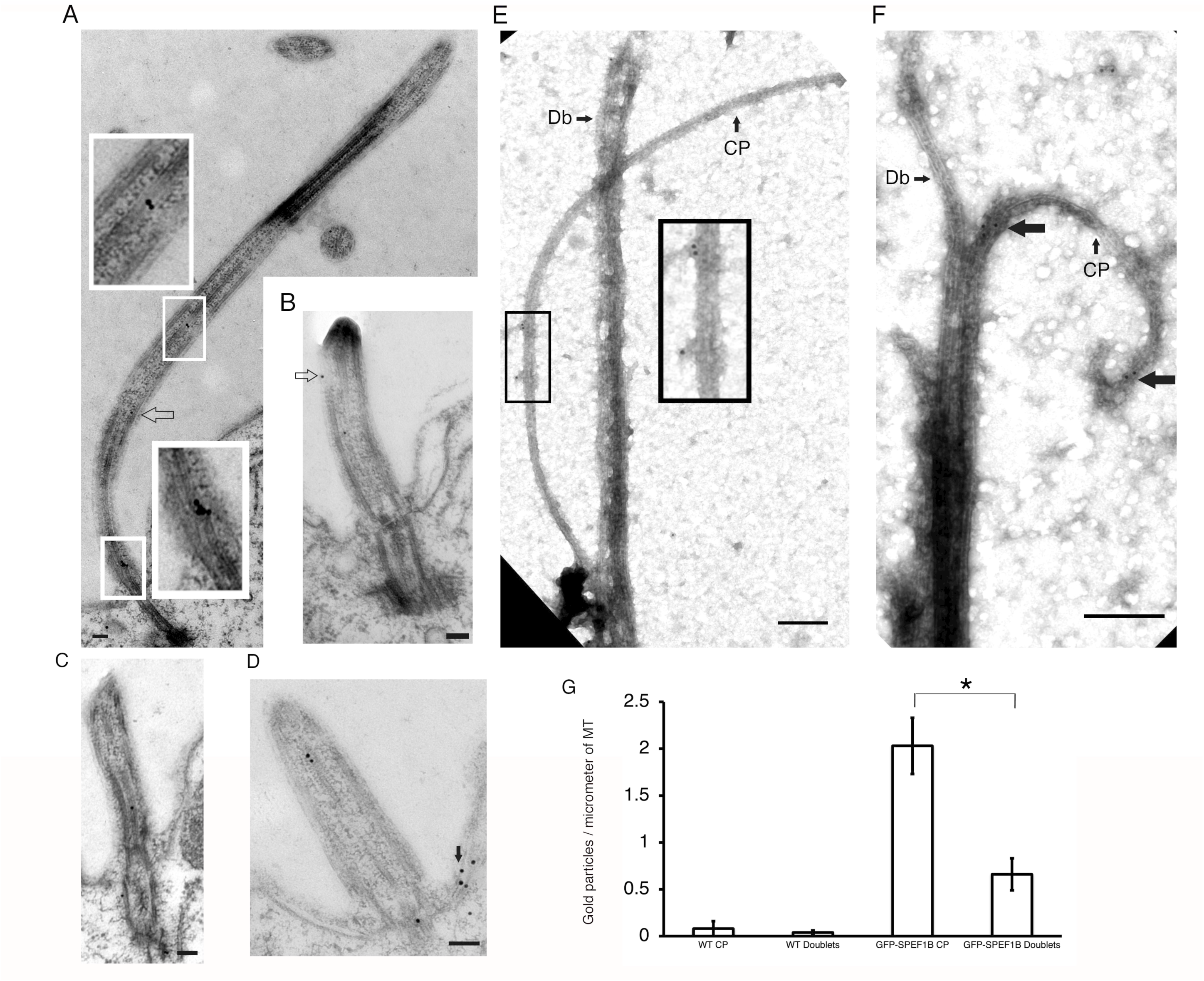
In the middle ciliary segment, SPEF1 preferentially localizes to the CP microtubules. (A-D) Post-embedding immunoelectron microscopy images of SPEF1B-mCherry-expressing cells (under native promoter). Inside the middle segment of cilia, SPEF1B is mainly present near the central microtubules (boxes in A show magnifications) and doublet microtubules (empty arrows in A and B). Outside of cilia, SPEF1B localized near the basal body and post-ciliary microtubules (black arrow in D). Scale bars = 100 nm. (E-F) Negatively stained ATP-reactivated axonemes isolated from the GFP-SPEF1B expressing cells (in the SPEF1AB-KO background, under *MTT1* promoter without induction with cadmium ions), stained with anti-GFP antibodies followed by secondary antibodies coupled to colloidal gold. Wild-type (not expressing a GFP fusion) axonemes were reactivated with ATP and probed with anti-GFP antibodies and anti-rabbit IgG-gold antibodies as negative control. GFP-SPEF1B is present on both central and doublet microtubules. The box in (E) shows a magnification of the CP. Larger arrows in (F) mark gold particles on the CP microtubules. Scale bars = 200 nm. (G) Quantification of the gold particle density on the central microtubules and doublets using whole mount images (such as those shown in panels E-F). Mean ± SE number of gold particles/micrometer of microtubule. WT CP: CP: 0.08 ± 0.08 (N = 13 axonemes); WT doublets: 0.04 ± 0.02 (N = 24); GFP-SPEF1B CP: 2.04 ± 0.03 (N = 16); GFP-SPEF1B doublets: 0.66 ± 0.17 (N = 22). *P = 0.002 (two tailed t test). Abbreviations: CP, central pair microtubules; Db-doublet.

### In cilia, SPEF1 particles are stationery and turnover slowly

Next, we analyzed mNeonGreen-SPEF1B in cilia in live (rescued SPEF1AB-KO) cells using TIRFM. In full length cilia, kymographs revealed a SPEF1B-enriched region at the distal end, as either one or multiple adjacent particles (Fig. 6A, kymographs of 4 cilia on the right, black arrowheads, Movie 6). A similar patchy pattern at the ciliary tips was observed by SR-SIM for both SPEF1A-GFP and mNeonGreen-SPEF1B (Fig. 4). Regardless of the number of segments, the tip-based SPEF1 particles retained their shape and remained stationary over a 10 sec observation time (Fig. 6A). In some cilia, there were particles outside of the tip region that also were stationary (Fig. 6A, cilia #2,4, red arrowheads). In cilium #1 (Fig. 6A, blue bracket) there is a nonuniformity in the SPEF1B signal across the middle segment consistent with diffusion of small SPEF1 particles. The kymographs (Fig. 6A) do not contain diagonal trajectories indicating an absence of IFT-mediated movements. Likely, inside cilia, SPEF1 distributes by diffusion and docks onto the microtubules, forming microdomains, with a prominent one across the distal compartment and smaller ones along the middle segment. To determine whether the SPEF1 particles, despite their fixed positions, undergo molecular exchange, we performed photobleaching of mNeonGreen-SPEF1B at the distal tip of a mature cilium. No recovery of mNeonGreen-SPEF1B signal over the period of 1 min was detected, indicating that SPEF1B turns over slowly (Fig. 6B and Movie 7).

**Figure 6.**
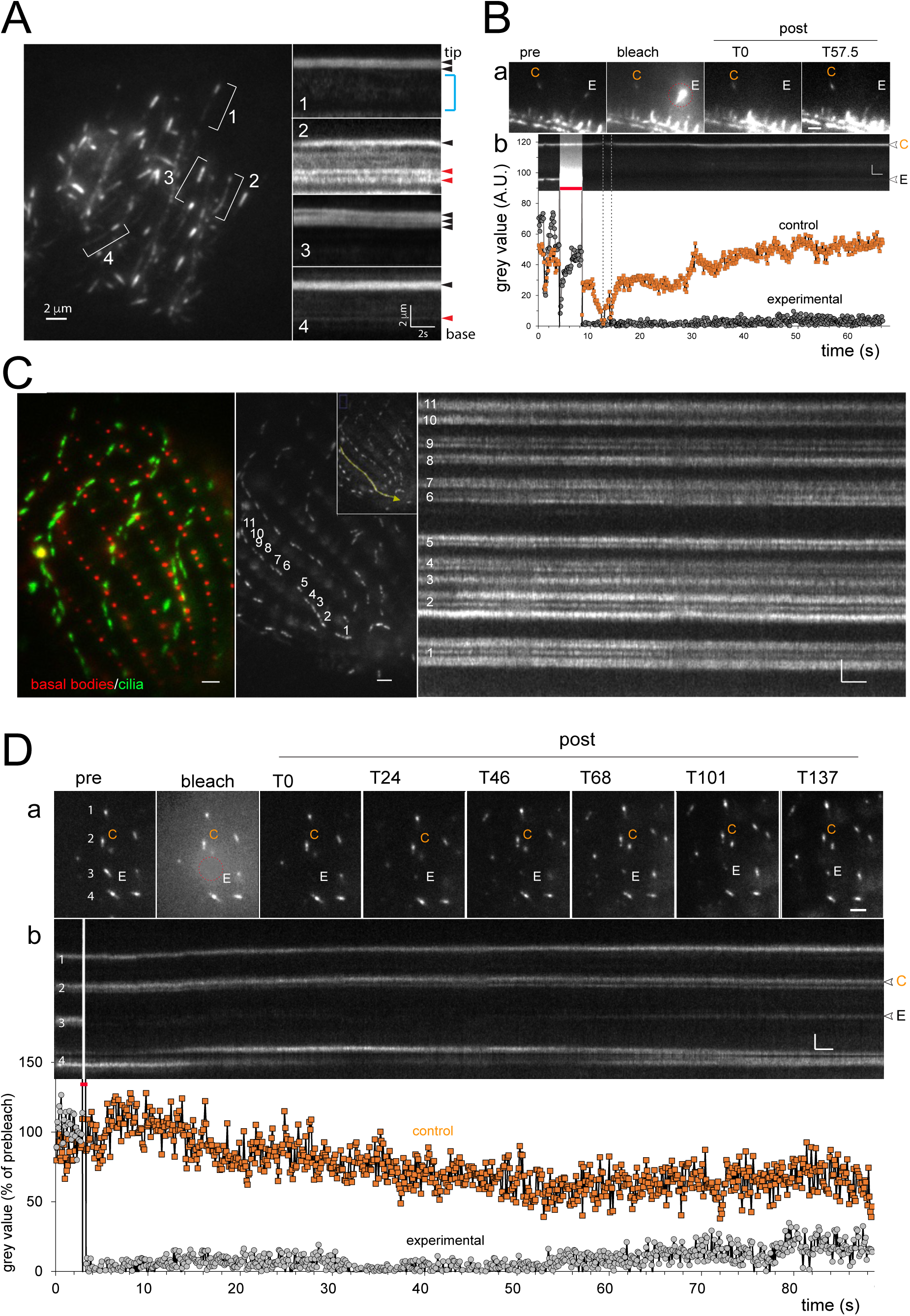
Ciliary SPEF1 particles are stationary and turnover slowly. TIRFM imaging of mNeonGreen-SPEF1B (expressed from an *MTT1*-driven transgene in the rescued SPEF1AB-KO strain, without induction by cadmium ions) in full-length (A-B) and growing (after deciliation) (C-D) cilia. All scale bars = 2 μm or 2 s. (A) Left panel: A vegetatively growing cell with 4 marked full-length cilia (#1-4) in which mNeonGreen-SPEF1B was imaged over time as documented in the kymographs on the right. Black and red arrowheads mark positions of SPEF1 particles at the cilium’s distal end or in the middle segment, respectively. In cilium #1 the blue bracket marks a weak scatterred signal likely representing diffusing SPEF1. Also see the corresponding Movie 6. (B) A FRAP experiment at the tip of a mature cilium. (a) Still images from a video showing two adjacent ciliary tips at various times before, during and after photobleaching of one tip with a focused laser beam (marked with E). The unbleached tip is a control (marked with C). (b) The kymograph shows the signal of mNeonGreen-SPEF1 at the two ciliary tips marked above, over time before, during (red period) and after photobleaching. The graph below shows a quantification of the signal intensities in the photobleached and untreated ciliary tip obtained suing above kymograph. See the corresponding Movie 7. (C) Imaging of growing cilia. Left panel: a merged image of a cilia-regenerating cell (at 1.5 hr post-deciliation), revealing mNeonGreen-SPEF1B at two focal planes: basal bodies (red) and ciliary tips (green). The cell is suspended in a small amount of medium which presses the imaged cilia against the coverglass. Middle panel: a single frame from a video at the level of ciliary tips. A kymograph analysis of 11 ciliary tips (numbered) was performed along the region marked with a yellow-green line and the resulting kymograph is shown on the right. See Movie 8. (D) A FRAP experiment at the tip of a regenerating cilium. (a) Still images of an area with multiple ciliary tips at various times before, during and after photobleaching of tips with a focused laser beam (marked with E). One of the unbleached tips is a control (C). (b) The kymograph documents the signal over time before, during (red period) and after photobleaching. The graph below shows a quantification of the signal intensities of the photobleached (E) and control (C) ciliary tip. Note that a limited recovery is apparent after 60 sec. See the corresponding Movie 9.

Next, we imaged SPEF1B in assembling cilia. Fig. 6C (based on Movie 8) shows a cilia-regenerating GFP-SPEF1B-expressing cell ∼2 hr after deciliation with multiple cilia, all having prominent tip signal. Consistently, there were between 2-4 resolvable SPEF1 particles per ciliary tip. In the assembling cilia, the pattern of SPEF1 was invariably composed of 2 or more particles After photobleaching, weak signal recovery was detectable after 1 min (Fig. 6D, Movie 9). Thus, at least in growing cilia, SPEF1 turns at a relatively slow rate (the structurally similar and tip-enriched EB1 recovered at ∼15% per min in the flagella of *Chlamydomonas* (Harris et al., 2016)). The presence of stationary foci and slow turnover indicate that a large pool of SPEF1 is bound to the microtubule lattice for a prolonged time, consistent with a role as microtubule stabilizer.

### SPEF1 loss alters the distribution of CP-associated MAPs

The two central microtubules (C1 and C2) are structurally and biochemically distinct (Carbajal-González et al., 2013; Gui et al., 2022; Han et al., 2022; Witman et al., 1972; Zhu et al., 2025). C1 and C2 are associated with distinct sets of MAPs that form meshworks either inside or on the surface of microtubules or are parts of large multi-protein projections (Cai et al., 2021; Carbajal-González et al., 2013; Chasey, 1969; Fu et al., 2019; Gui et al., 2022; Han et al., 2022; Linck et al., 1981; Zhu et al., 2025). To evaluate how the loss of SPEF1 affects the CP complexes, we tagged two well-studied CP MAPs, PF16 and hydin (Hyd1), with GFP (by engineering the corresponding native loci) in the wild-type and SPEF1AB-KO backgrounds. PF16 oligomerizes into a spiral on the C1 microtubule surface and provides a base for attachment of the C1 projections (Gui et al., 2022; Han et al., 2022; Smith and Lefebvre, 1996; Zhu et al., 2025). Hydin binds at multiple points around C2 and extends to C1, contributing to the bridge that joins C1 and C2 (Gui et al., 2022; Han et al., 2022; Lechtreck and Witman, 2007; Zhu et al., 2025). Polyglycylation of tubulin is a conserved post-translational modification that is abundant on axonemal microtubules (Levilliers et al., 1995; Redeker et al., 1994). In *Tetrahymena*, anti-polyglycylated tubulin (polyG) antibodies detect this PTM on the outer but not on CP microtubules (Wloga et al., 2009). In agreement, using SR-SIM, the signal of tubulin polyglycylation formed two parallel lines. As expected, in the wild-type background, PF16-GFP and hydin-GFP signals formed single lines, flanked by the polyglycylation lines (Fig. 7A,B top panels, 7C,D left panels). In the wild-type cilia, the PF16-GFP signal was uniform from the base to just below the distal tip (Fig. 7A,C), whereas in the SPEF1AB-KO cilia, the PF16-GFP signal was severely disturbed (Fig. 7A,C); 25% of cilia lacked the PF16-GFP line entirely and 46% of cilia had either shorter or fragmented PF16-GFP lines (Fig. 7A,E). In the wild type, the hydin signal was also a single uninterrupted line between the base and the tip (Fig. 7B,D). The defects in the hydin organization in the SPEF1AB-KO cells were less severe. All SPEF1AB-KO cilia examined had hydin signals but in ∼15% of them, there were gaps in the hydin-GFP line (Fig. 7B,D,F); sometimes the ends of the hydin-GFP lines on the two sides of the gap seemed out of register suggesting that they flank a small microtubule gap (Fig. 7D, yellow arrows). As described above, TEM revealed that cross-sections lacking a single microtubule were more frequent as compared to those lacking both microtubules and longitudinal sections showed an apparent loss of the proximal portion of C1. The strong effect on the distribution of PF16, a principal component of the C1 surface, supports the earlier conclusion that C1 is more sensitive to the loss of SPEF1. Likely, the pattern of hydin is less affected because most of the hydin binding interface is on C2 (Lechtreck and Witman, 2007; Zhu et al., 2025).

**Figure 7.**
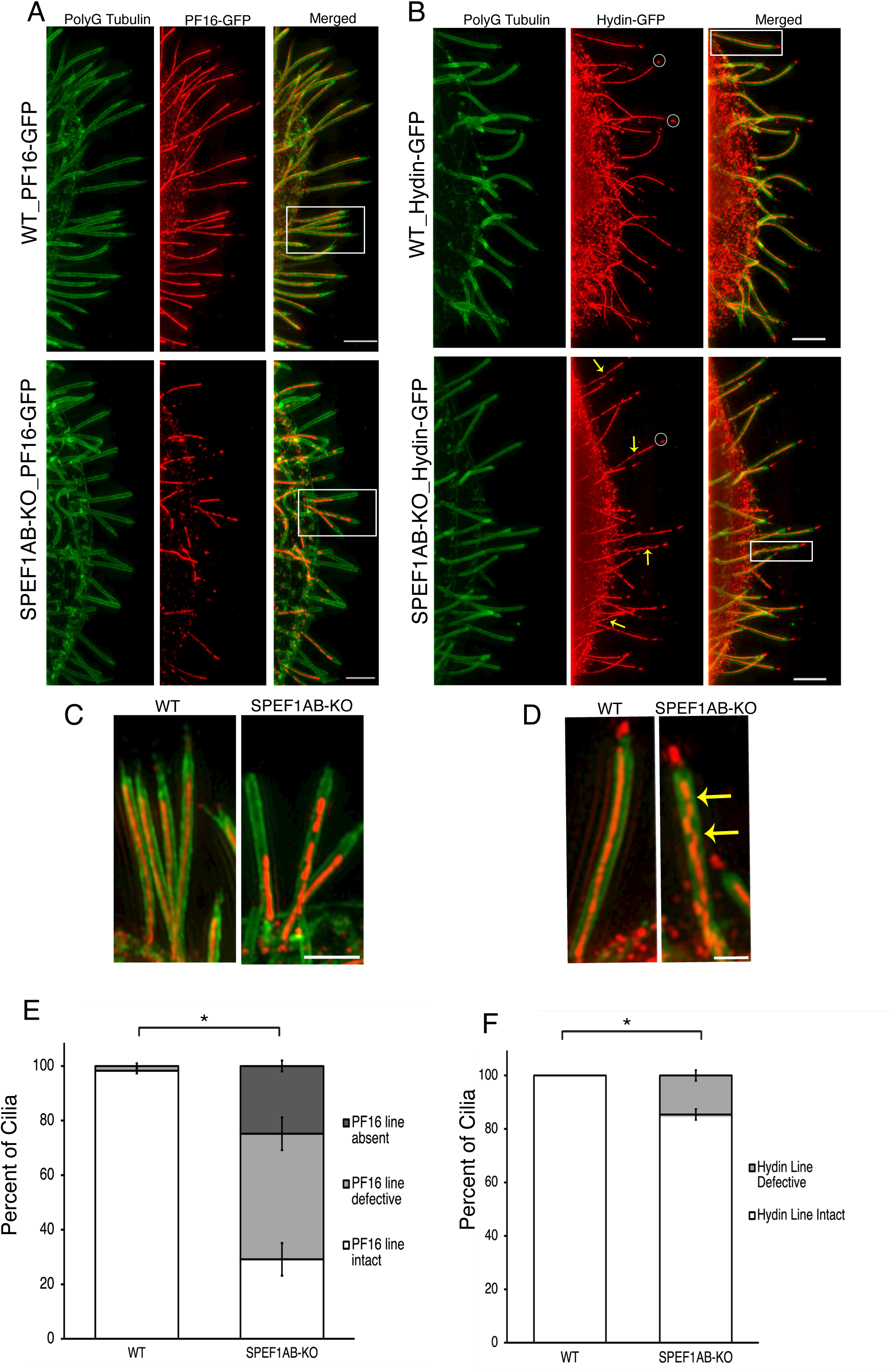
Loss of SPEF1 damages the CP microtubule complexes. (A-D) SR-SIM images of wild-type and SPEF1AB-KO cells expressing either PF16-GFP (A,C) or hydin-GFP (B,D). Cells were labeled antibodies against polyglycylated tubulin (green) and anti-GFP antibodies to detect either PF16-GFP (red) or hydin-GFP (red). In B, the white circles at the tips of cilia shows non-specific labeling by anti-GFP antibodies (based on the negative controls). Scale bars = 3 μm. (C,D) Areas marked by boxes in A and B shown at a higher magnification. In D, the pattern of hydin-GFP indicates that the CP complex has internal gaps and the CP complex ends that surround the gap are displaced (yellow arrows). Scale bars: 1.5 μm in C and 0.75 μm in D. (E) Quantification of the patterns of PF16-GFP lines. Percentages of cilia with an intact, fragmented or absent PF16-GFP line per cell. WT: N = 8 cells, N = 171 cilia; SPEF1AB-KO: N = 10 cells, N = 159 cilia. Error bars are standard errors. *P < 0.0001 (two tailed t test) for each category scored. (F) Quantification of the hydin-GFP line patterns. Percentages of cilia with an intact or defective hydin-GFP line per cell. WT: N = 8 cells, N = 102 cilia; SPEF1AB-KO: N = 11 cells, N167 cilia. Error bars represent standard errors. *P = 0.0003 (two tailed t test) for each of the 3 categories scored.

### Cryo-electron tomography reveals defects in the CP tubulin lattice and a loss of CP projections

To examine defects in the CP at higher resolution, we performed cryo-ET of the SPEF1AB-KO and wild-type cilia. The longitudinal and cross sections of the mutant axonemes revealed CPs lacking portions of the microtubule wall (Fig. 8B,D black arrows, compare to 8A) and CP segments devoid of lateral projections, probably on C1 (Fig. 8C top, blue bracket). The cross-section of the CP region lacking projections on C1 showed an abnormal proximity of C1 and C2 (Fig. 8C bottom, blue bracket compare to Fig. 8A bottom). Unexpectedly, there were abnormal densities outside of the A-tubule of the SPEF1AB-KO doublet microtubules, close to the seam position (between protofilaments A9 and A10) (Fig. 8B,D red arrows). Finally, in the SPEF1AB-KO axonemes that retained the plasma membrane (likely due to incomplete demembranation by the detergent), there was an abnormally dense ciliary matrix, suggesting that the mutant cilia accumulate materials (Fig. 8E yellow arrowheads). To conclude, the loss of SPEF1 damages the walls of CP microtubules and generates regions that lack projections but have seemingly intact microtubules. Thus at least some of the gaps in the distribution of PF16 and hydin seen in SR-SIM may represent segments missing MAPs on otherwise intact or perhaps more likely mildly damaged CP microtubules.

**Figure 8.**
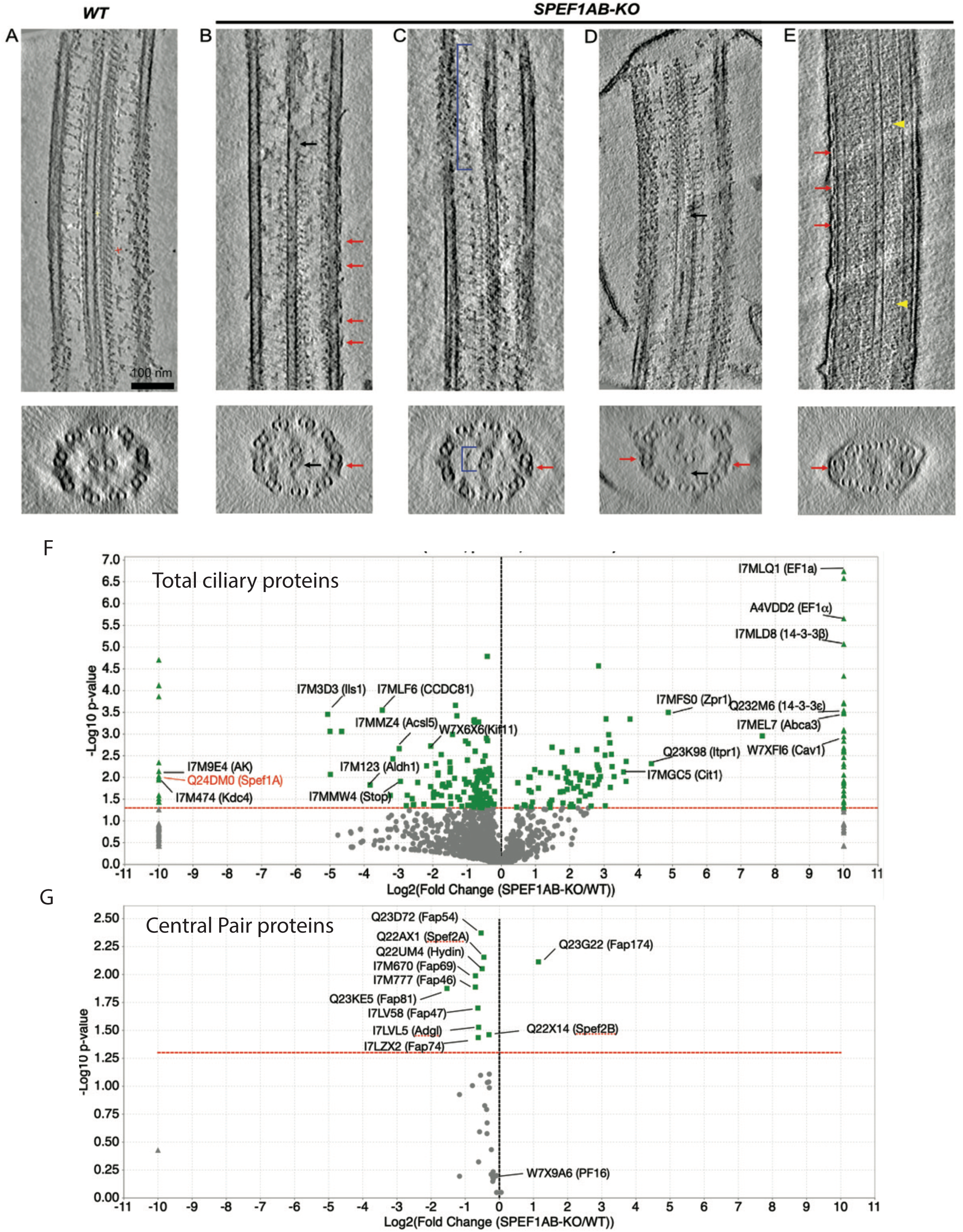
Loss of SPEF1 damages the lattice of central microtubules and alters the proteome of cilia. (A-E). Cryo-ET images of wild-type (A) and SPEF1AB-KO (B-E) axonemes within the middle segment. (A) Longitudinal and cross-sectional views of the wild-type (A) and SPEF1AB-KO axonemes (B-E). In (E), the axoneme has an intact membrane likely due to incomplete detergent solubilization. Black arrows indicate damaged walls of central pair microtubules. Red arrows mark abnormal densities on the doublet microtubules. Yellow arrowheads point to dense ciliary matrix inside the SPEF1AB-KO cilia. In C, the blue bracket marks the CP region lacking the lateral projections. In the cross-section within the middle segment shown below, the two central pair microtubules are positioned abnormally close to each other, assuming the distance the distance that the CP microtubules are separated by in the distal tip region. (F-G). Volcano plots show the relative levels of proteins detected by mass spectrometry between the wild-type and SPEF1AB-KO cilia. All detected proteins are analyzed in (F) and known CP subunits are analyzed in (G).

### The loss of SPEF1 loss impacts the ciliary proteome

The unexpected abnormalities in the SPEF1AB-KO cilia revealed by cryo-ET (dense cytoplasm and abnormal projections) prompted us to explore the proteomes of SPEF1AB-KO and wild-type cilia using mass spectrometry. Among a total of 2794 proteins detected, 151 were significantly depleted and 124 proteins were enriched in the SPEF1AB-KO versus wild-type cilia, respectively (Fig. 8F, Table S3). As expected, SPEF1A was absent in SPEF1AB-KO cilia (SPEF1B was not detected in both the wild-type and SPEF1AB-KO cilia). Among the evolutionarily conserved proteins not detected in the SPEF1AB-KO cilia but present in the wild-cilia was an arginine kinase (AK, UniProtID: I7M9E4). AKs are associated with the ciliary membrane and may regulate the levels of ATP for dynein-mediated ciliary motility, acting as a phosphagen shuttle (Kutomi et al., 2012; Noguchi et al., 2001; Watts and Bannister, 1970; Yano et al., 2020). AK levels were also altered by deletions of radial spoke proteins (Bicka et al., 2025). Abnormal AK levels indicate an imbalance in the ciliary energy metabolism, perhaps secondary to the reduced ciliary motility. SPEF1 is highly enriched within the ciliary distal tip compartment where it contributes to the mutual alignment of C1 and C2 (Legal et al., 2025), and therefore its loss may affect the levels of other distal tip-resident proteins. Among the highly depleted (>8-fold) proteins, there was Ccdc81, a ciliary distal tip-enriched protein, discovered based on the dependence of its ciliary presence on another conserved ciliary tip protein, Fap256/Cep104 (Legal et al., 2023; Louka et al., 2018; Tammana et al., 2013). In addition, the SPEF1AB-KO cells had reduced level of a STOP domain protein (UniProtID: I7MMW4), whose presence in cilia also depends on Fap256/Cep104, (Legal et al., 2023). Several known tip proteins were detected (Fap256A and Fap256B, Armc9A, Tlp1, Tlp2, C2d1, Spike1, Spike2 and Fap213 (Legal et al., 2025; Louka et al., 2018) but their levels were unaltered. Thus, in agreement with the ultrastructural assessment ( (Legal et al., 2025) and this study), the proteomic data indicate that despite its strong enrichment, SPEF1 has a minor role in distal segment organization.

As documented above, the loss of SPEF1 dramatically affects the organization of the CP microtubules and distribution of some CP MAPs within the middle segment. Among the 39 detected CP proteins of the middle segment (Fig. 8G; Table S4), 9 were depleted in SPEF1AB-KO, including Fap74 and Fap81 that are located on the surface of C1 and connect to the bases of C1a/e/c and C1d projections, respectively (Brown et al., 2012; Fu et al., 2019; Gui et al., 2022; Han et al., 2022; Zhao et al., 2019), components of the C1d projection: Fap46 and Fap54 (Brown et al., 2012; Gui et al., 2022; Han et al., 2022), components of C1b: Fap69, Adgl, two paralogs of Spef2/Cpc1 (Cai et al., 2021; Gui et al., 2022; Han et al., 2022; Joachimiak et al., 2021). Also depleted in SPEF1AB-KO cilia were also Fap47 and hydin, which are constituents of the C1-C2 bridge and have binding interphases on both C1 and C2 (Gui et al., 2022; Han et al., 2022; Zhu et al., 2025). On the other hand, a component of C1b, Fap174 (Cai et al., 2021; Gui et al., 2022; Joachimiak et al., 2021) was enriched. The levels of 4 detected proteins detected known to be associated with C2 (Fap65, Fap70, Klp1, Fap213) were unchanged (Table S4). Thus, all affected proteins are attached to C1, providing further evidence that SPEF1 plays a particularly important role in C1 integrity. Surprisingly, the level of the major C1 outer surface protein PF16 (Gui et al., 2022; Han et al., 2022; Zhu et al., 2025) that was missing or fragmented in the permeabilized/fixed SPEF1AB-KO cilia (Fig. 7A,E) was normal (Table S4). Likely, unassembled PF16 remains inside the SPEF1AB-KO cilia and contributes to the increased density of the ciliary matrix (Fig. 8E). Our observations agree with the presence of unassembled CP proteins in the 9+0 mutant cilia of *Chlamydomonas* (Lechtreck et al., 2013). Unexpectedly, 24 proteins were present in the SPEF1AB-KO and not in the wild-type cilia (Fig. 8F, Table S3), including two orthologs of the eukaryotic translation elongation factor 1 alpha, EF1a (UniProtID: A4VDD2 and I7MLQ1). EF1a was purified from *Tetrahymena* cilia and detected on axonemal microtubules by immunoelectron microscopy (Numata et al., 2000). On the other hand, EF1a accumulates in cilia of *Chlamydomonas reinhardtii* mutants lacking CEP290, a component of the transition zone (Craige et al., 2010). Thus, the accumulation of EF1a could reflect a lack of proper gating between cilia and the cell body compartment. To further explore the trends in the protein composition induced by SPEF1 loss, we focused on 3 groups: microtubule inner proteins (MIPs) of outer doublets, subunits of radial spokes (RSPs) and IFT proteins. The MIPs, which form a reinforcing meshwork inside the lumen of the doublet microtubules (Fabritius et al., 2021; Howard-Till et al., 2025; Kubo et al., 2023a; Li et al., 2022; Stoddard et al., 2018), were nearly unaffected; among a total 41 detected, one was slightly elevated (Fig. S6, Table S5). Radial spokes interact with the CP projections, mediating mechanosensory signals that control the dynein activity (reviewed in (Zhu et al., 2017)). Among the 39 RSPs (Bicka et al., 2025; Bicka et al., 2022) detected, 3 were mildly depleted (Fig. S6, Table S6). Interestingly, among the 47 IFT proteins detected (components of IFT trains, motors and cargo adapters including the BBS complex (Lacey and Pigino, 2025; Lechtreck, 2022)), 7 were depleted over 2-fold and none were enriched in the SPEF1AB-KO cilia (Fig. S6, Table S7). Curiously, 5 of the depleted IFT proteins are components of IFT train subcomplex A, which is particularly important for the retrograde IFT (Piperno et al., 1998): IFT122, IFT139, IFT140, IFT121 and IFT144. A preferential depletion of the IFT complex A was reported in *Tetrahymena* mutants lacking subunits forming the C1b/f projection (Joachimiak et al., 2021). On the other hand, in the *Chlamydomonas* mutant 9+0 cilia, several IFT proteins (both A and B) are elevated (Lechtreck et al., 2013). Taking together the data from two ciliated species, CP deficiencies change the levels of IFT proteins. Abnormal IFT is likely related to reduced ciliogenesis observed in the CP mutants but the cause-effect relationship is unclear.

### SPEF1 is required for proper assembly of the CP complexes

The defects in the CP structure caused by SPEF1 loss may arise during ciliary assembly or may reflect a lack of resistance to mechanical stress when cilia beat vigorously. To test whether SPEF1 is important for CP assembly, we first examined how the C1 complex normally forms in the wild-type, cilia-regenerating cells, using PF16-GFP as a reporter. At 30 min post-deciliation, most cilia were short, (below ∼1.5 μm) and ∼25% had PF16-GFP as a single complete line, usually at a small distance from the cilium base (Fig. 9A). In longer cilia observed at 60 min, the PF16-GFP signal pattern was variable: absent, consisting of a single or multiple segments, or forming a complete line of PF16-GFP (Fig. 9A). A similar variable pattern of PF16-GFP was seen at 90 min although the proportion of cilia with a single line pattern had increased. At 120 min, most cilia had as a single complete line (Fig. 9A). PF16 is an important MAP that polymerizes into a spiral on the surface of C1 and contributes to docking of C1 projections (Gui et al., 2022; Han et al., 2022). Although we did not directly image the assembly of C1 microtubule, it is evident that maturation of the C1 microtubule complex lags behind the axoneme elongation, which is driven by polymerization of axonemal microtubules. Furthermore, it appears that the complexes covering the surface of C1 are seeded at multiple sites along the cilium length and later integrate into one line. Moreover, the patchy pattern of PF16-GFP in the assembling wild-type cilia resembles that of the mature SPEF1AB-KO cilia. Thus, the CP defects observed in the mature cilia lacking SPEF1 may result from an incomplete CP complex assembly. To explore this possibility further, we compared the pattern of PF16-GFP during cilia regeneration between the wild-type and SPEF1AB-KO backgrounds. In the wild type, 16% of cilia had an “immature” patchy pattern of PF16-GFP at 3 hr, which decreased to 3-4% at 5 and 7 hr of regeneration (Fig. S7). In contrast, in the SPEF1AB-KO cells, 41% of cilia had a patchy PF16-GFP pattern, followed by 53 and 65% at 5 and 7 hr, respectively (Fig. S7; cilia lacking PF16-GFP entirely were not included in this quantification). Thus, the defects observed in the absence of SPEF1 are already apparent during ciliary assembly.

**Figure 9.**
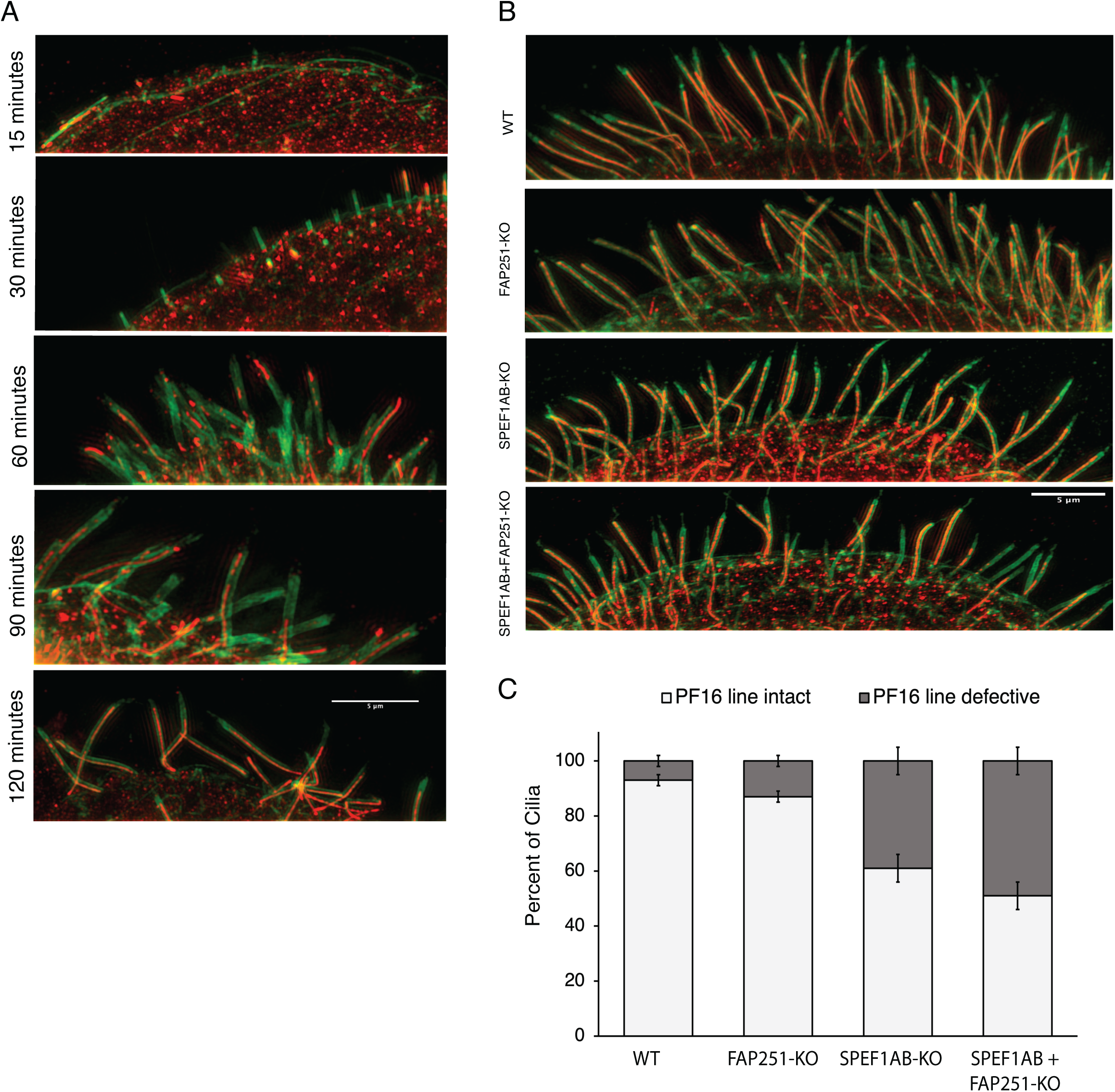
The CP defects caused by the loss of SPEF1 are apparent during cilia assembly and are not rescued by genetic reduction of ciliary motility. (A) SR-SIM images of wild-type cells expressing PF16-GFP, fixed at various time points after pH shock deciliation, labeled by anti-polyglycylated tubulin antibodies (green) and anti-GFP antibodies to detect PF16-GFP (red). Scale bar = 5 μm. (B) Images of cells expressing PF16-GFP (red) and labeled by the anti-polyglycylated tubulin antibodies (green) with the following genotypes: WT, FAP251-KO, SPEF1AB-KO, SPEF1AB + FAP251-KO (a triple knockout strain). *Tetrahymena cells* lacking FAP251, a component of RS3 base, have a severely reduced rate of ciliary beating that result in partial paralysis (Urbanska et al., 2015). (C) The graph quantifies the frequency of defects (cilia with fragmented or absent PF16-GFP lines) as a function of the genotype.

While the observations made so far suggest that SPEF1 is important for the CP assembly, a possibility remains that SPEF1 stabilizes the CP against mechanical stress generated by axoneme bending as even the shortest cilia are motile (Rosenbaum and Carlson, 1969). To test whether the CP defects are enhanced by ciliary motility, we deciliated wild-type and SPEF1AB-KO cells (expressing PF16-GFP) and allowed them to regenerate cilia in the presence of 500 μM nickel chloride, to inhibit the ATPase activity of the inner dynein arms (Bayless et al., 2012; Larsen and Satir, 1991; Meehl et al., 2016). Nearly all scored wild-type cells assembled a single line C1 in the presence of NiCl_2_ (Fig. S7A). However, NiCl_2_ did not rescue the C1 assembly defects in the SPEF1AB-KO cilia (Fig. S7A,B). Finally, we tested whether the C1 defects can be rescued by reducing ciliary motility using a genetic tool, namely by deleting Fap251, a conserved component of the RS3 radial spoke, whose loss causes ∼80% reduction in the cell swimming velocity in *Tetrahymena* (Urbanska et al., 2015). Similar to the chemical treatment, genetic reduction of ciliary motility did not rescue the C1 defects in the SPEF1AB-KO cilia (Fig. 9B,C). It appears therefore that the mechanical stress caused by ciliary motion is not a major driver of the structural defects in the CP and that SPEF1 may play a direct role in the assembly of CP microtubule complexes.

### SPEF1 coevolves with the central pair apparatus

SPEF1 associates with both ciliary and non-ciliary microtubules in *Tetrahymena*, *Trypanosoma* and animal cells (Chan et al., 2005; Kim et al., 2018; Legal et al., 2025; Pham et al., 2022; Tapia and Hecht, 2022; Tapia et al., 2025; Werner et al., 2014; Zheng et al., 2019). In light of its broad subcellular distribution, it is surprising that the defects caused by the loss of SPEF1 are so far apparent only in the central microtubules ((Wu et al., 2025; Zheng et al., 2019) and this study). We therefore examined the phylogenetic pattern of SPEF1 homologs identified by reciprocal pBLAST searches (E value <1e-50) (Table 1). SPEF1 homologs were found only in ciliated species. However, SPEF1 was absent in the predicted proteomes of two ciliated species that lack the central apparatus, namely *C. elegans* (Doroquez et al., 2014) and the diatom *Thalassiosira pseudonana* (Idei et al., 2013). It therefore appears that SPEF1 has co-evolved with the CP apparatus of motile cilia, which fits with its important role for CP organization.

**Table 1.** Phylogenetic distribution of SPEF1, cilia and the central microtubule apparatus across diverse ciliated and non-ciliated lineages. SPEF1 orthologs were identified by reciprocal pBLAST searches using human and *Tetrahymena* sequences (E value <1e-50).

| Species | Cilia | Central Pair Apparatus | SPEF1 |
| --- | --- | --- | --- |
| *Homo sapiens* | ✓ | ✓ | ✓ |
| *Danio rerio* | ✓ | ✓ | ✓ |
| *Ciona intestinalis* | ✓ | ✓ | ✓ |
| *Caenorhabditis elegans* | ✓ |  |  |
| *Drosophila melanogaster* | ✓ | ✓ | ✓ |
| *Hydra magnipapillata* | ✓ | ✓ | ✓ |
| *Nematostella vectensis* | ✓ | ✓ | ✓ |
| *Monosiga brevicollis* | ✓ | ✓ | ✓ |
| *Ustilago maydis* |  |  |  |
| *Schizosaccharomyces pombe* |  |  |  |
| *Saccharomyces cerevisiae* |  |  |  |
| *Aspergillus fumigatus* |  |  |  |
| *Batrachochytrium dendrobatidis* | ✓ | ✓ | ✓ |
| *Encephalitozoon cuniculi* |  |  |  |
| *Entamoeba histolytica* |  |  |  |
| *Dictyostelium discoideum* |  |  |  |
| *Ostreococcus tauri* |  |  |  |
| *Chlamydomonas reinhardtii* | ✓ | ✓ | ✓ |
| *Arabidopsis thaliana* |  |  |  |
| *Physcomitrella patens* | ✓ | ✓ | ✓ |
| *Cyanidioschyzon merolae* |  |  |  |
| *Plasmodium falciparum* | ✓ | ✓ | ✓ |
| *Tetrahymena thermophila* | ✓ | ✓ | ✓ |
| *Thalassiosira pseudonana* | ✓ |  |  |
| *Trichomonas vaginalis* | ✓ | ✓ | ✓ |
| *Trypanosoma cruzi* | ✓ | ✓ | ✓ |
| *Naegleria gruberi* | ✓ | ✓ | ✓ |

## Discussion

The phylogenetic distribution suggests that the emergence of SPEF1 was an important step during evolution of motile 9+2 cilia. Over time, SPEF1 may have acquired functions on microtubule rootlets associated with the basal bodies (Pham et al., 2022), which were likely already present in the most recent ancestor of modern eukaryotes (Yubuki et al., 2016). However, we show here that in *Tetrahymena*, despite SPEF1 being present on diverse types of microtubules, its loss detectably affects only the CP microtubules (with the caveat that we did not examine the non-ciliary microtubules with the same level of scrutiny as the axoneme). In the cilium, SPEF1 is typically concentrated within the ciliary distal tip region where it binds along the seam of the C1 and C2 (Legal et al., 2025). Surprisingly, despite its distal enrichment, in *Tetrahymena*, the loss of SPEF1 dramatically disorganized the CP in the middle segment, which comprises most of the cilium length. The absence of SPEF1 damaged the walls of the CP microtubules and led to a loss of lateral projections, the latter of which is likely due to underlying defects in the tubulin lattice. Clearly, both CP microtubules are affected by the loss of SPEF1, as indicated by the presence of 9+0 axoneme cross-sections ((Wu et al., 2025; Zheng et al., 2019) and this study), which agrees with the presence of SPEF1 along the seam of both C1 and C2 in the distal tip region in *Tetrahymena* cilia (Legal et al., 2025). However, at least in *Tetrahymena*, the C1 complex is more sensitive to the loss of SPEF1 than C2 complex. Our observations also reveal that without SPEF1, the CP microtubules of the middle segment acquire assembly defects. In contrast, in the distal segment, SPEF1 appears to play a subtle role by aligning the two CP microtubules, possibly through its ability to form a dimer that cross-links C1 and C2 (Legal et al., 2025). In *Tetrahymena*, the distal enrichment of SPEF1 is established early during cilium outgrowth ((Legal et al., 2025) and this study). It is therefore possible that the widespread middle segment defects are a catastrophic consequence of relatively small positional shifts within the CP at the ciliary tip during cilium assembly (Legal et al., 2025). However, we show here that SPEF1 is present along the CP microtubules in the middle segment, where it forms relatively immobile particles, opening a possibility that SPEF1 has a CP microtubule-stabilizing along the entire cilium shaft.

One possibility we have explored here is that SPEF1 protects the CP microtubules against mechanical stress during the ciliary beat cycle. Although the widespread CP defects were not rescued by reducing ciliary motility, a role for SPEF1 in mechanical stability cannot be excluded. The CP microtubules are under strain that is intrinsic to the CP apparatus. When the CP is extruded from the axoneme *in vitro*, it adopts a helical shape (Omoto et al., 1999; Omoto and Kung, 1979; Omoto and Kung, 1980). Both *in vivo* and in an extruded CP, C1 is positioned on the convex side (Bernstein et al., 1994; Lechtreck and Witman, 2007; Mitchell, 2003). Beams under bending exhibit lower tensile strength compared to the compressive strength, and therefore defects are more likely to occur on the convex side (Ashby, 2011). Furthermore, the CP microtubules are twisted around each other along the longitudinal axis (Carbajal-González et al., 2013; Goodenough and Heuser, 1985; Han et al., 2022). The differences in the MAP composition between C1 and C2 may further contribute to the differential dependence on structural reinforcement by SPEF1. Within the middle segment, the C2 MAP complexes (including projections and microtubule surface proteins) follow a 16 nm periodicity, while a subset of MAP complexes on C1 are repeated every 32 nm (Carbajal-González et al., 2013; Gui et al., 2022; Han et al., 2022; Zhu et al., 2025). Moreover, the seam of C2 is decorated by a complex of PF20 and FAP178 with an 8 nm periodicity (Gui et al., 2022; Han et al., 2022; Zhu et al., 2025) and could therefore substitute for the SPEF1 seam reinforcement specifically on C2. Overall, C2 has a denser MAP coverage, which could explain the reduced dependence of C2 on stabilization by SPEF1. Finally, in *Tetrahymena*, the end of C1 is embedded inside the axosome, while the end of C2 is not (Hausmann and Fischer-Defoy, 1978). Thus, C2 may have more conformational freedom to release the tension that accumulates as the two microtubules are subjected to bending and twisting.

Alternatively, SPEF1 may be important for polymerization of tubulin to form the lattice of CP microtubules. We show here that in the assembling wild-type cilia, the principal C1 surface MAP, PF16 (Gui et al., 2022; Han et al., 2022), is assembled in patches that with time integrate into a single line. Thus, it appears that at least some CP MAPs are seeded at multiple locations along the length of elongating CP microtubules. Intriguingly, at least under certain conditions, the central microtubules themselves can be nucleated away from the cilium base. In mating dikaryons of *Chlamydomonas reinhardtii* composed of a wild-type and a 9+0 mutant, during CP repair in the mutant flagella, segments containing both tubulin and hydin first appear away from the ciliary base, often near the cilium distal end (Lechtreck et al., 2013). Under normal conditions, in the very short assembling cilia, the ends of the CP microtubules are already connected to the transition zone (Lechtreck et al., 2013). It remains possible, that the CP assembly is initiated away from the cilium base (possibly by microtubule seeds as suggested by Liu and colleagues (Liu et al., 2021)) during a very brief period that may be difficult to capture by electron microscopy, followed by docking of the minus end of the nascent CP at its position immediately proximal to the transition zone. Reinforcement of the microtubule seam by SPEF1 may be important for the unique mode of CP assembly. Alternatively, a recent study reports that SPEF1 dimers form condensates with tubulin *in vitro* (Ren et al., 2025), raising the possibility that SPEF1 concentrates tubulin building blocks for the CP assembly.

To summarize, we show that SPEF1 is broadly distributed throughout the *Tetrahymena* cell but plays a critical role in organizing the CP microtubules within the middle segment of cilia. In addition to its high level in the ciliary distal segment, SPEF1 is present along the ciliary middle segment and is enriched on the central relative to the outer microtubules. We propose that SPEF1 reinforces the microtubule lattice in the middle segment and that this activity is critical for the CP assembly.

## Materials and Methods

### Tetrahymena thermophila strains and culture

*Tetrahymena thermophila* strains were grown at 30°C in SPP medium (Gorovsky, 1973) supplemented with antibiotics (SPPA). CU428 strain was used as a wild type (RRID: TSC_SD00178, *Tetrahymena* Stock Center, Washington University in St. Louis, MO).

### Generation of germline knockouts

Using homologous DNA recombination in *Tetrahymena thermophila,* we disrupted the coding regions of *SPEF1A* (*TTHERM_00939230*) and *SPEF1B* (*TTHERM_00161270*) background (Legal et al., 2025). To evaluate the CP MAP distributions, we edited genes to encode C-terminal GFP fusions: *PF16* (*TTHERM_000157929*) and hydin (*HYD1/TTHERM_00551040*) using plasmid targeting fragments with a linked *neo5* marker (Busch et al., 2010). *PF16* and *HYD1* were edited in multiple backgrounds: wild-type (strain CU428), SPEF1AB-KO and FAP251-KO (Urbanska et al., 2015) and in a triple knockout strain SPEF1AB-KO;FAP251-KO, by biolistic transformation and selection with 100 μg/ml paromomycin. The transgene copy number was increased by phenotypic assortment by growing cells in an increasing concentration of paromomycin up to 3 mg/ml.

### Phenotypic studies in *Tetrahymena*

To measure the swimming velocity, cells were grown to a density of 2 x 10^5^ cells/ml and imaged in dark-field on a Nikon Eclipse 55i microscope using a 40X Nikon Plan objective. Images were captured using 1 sec exposure on a Zeiss AxioCam ERc 5s camera. Observations of cilia waveform and quantification of cilia beat frequency were performed as described (Funfak et al., 2015; Stoddard et al., 2018). A highly concentrated cell suspension was loaded into a microfluidic channel (33 μm in height and 50 μm) using a controllable pressure source (Fluigent). The pressure in the channel was equilibrated to prevent residual flow, by adjusting the pressures in the inlet and outlet of the channel. Forward-swimming cells were imaged using a 90X objective by differential interference contrast on an inverted microscope (Nikon TE2000). Videos were recorded at 2000 frames per second using a Photron 1024PCI camera. Matlab and CR Toolbox were used to mark the cell body outline and the zone of beating cilia in the first frame of each movie. The moving cell was tracked by repositioning the cell/cilia boundaries every 30-50 frames. The cilia zone was digitally unwrapped producing a rectangular video of beating cilia in which all ciliary bases were aligned horizontally. Fiji/ImageJ was used to create a space-time diagram (chronograph) of beating cilia by plotting the greyscale at a fixed position away from the cell body, over time. Ciliary beat frequency was quantified by measuring the distances (on the y-axis) between adjacent diagonal lines corresponding to the same cilium. To measure the rate of phagocytosis, cells were grown to a density of 2 x 10^5^ cells/ml and fed with India ink (0.1 %) for 30 min. Cells were fixed with 2% paraformaldehyde and the number of ink-filled food vacuoles per cell was scored using bright field microscopy. The multiplication rates were determined by growing cells in 5 ml cultures (starting at 1 x 10^4^ cells/ml) and the cell density was measured at multiple time points. Confocal images of cells stained with anti-polyglycylation and anti-centrin antibodies were used for quantifications of the cilia number, length and rate of assembly (after deciliation (Calzone and Gorovsky, 1982)). Quantifications were done using the methods as described in (Dave et al., 2009). To inhibit cell motility, cells were treated with 500 μM NiCl_2_ (Jiang et al., 2015).

### Evaluation of *spef1* in zebrafish

Tupfel Longfin (TL) zebrafish embryos were maintained at 28.5°C in E3 medium (NaCl 0.287 g/L, KCl 0.0132 g/L, CaCl₂·2H₂O 0.0479 g/L, MgCl₂·6H₂O 0.0807 g/L, 0.00005% methylene blue, pH 7.2). Antisense morpholino oligonucleotides (made by Gene Tools LLC) were designed to target the intron 1-exon 2 splice site of *spef1* (5-AGCCATAACTGCTTGCACAAATCGA-3’), to disrupt splicing of *spef1*, leading to aberrant transcripts predicted to introduce premature termination codons and trigger nonsense-mediated decay. Morpholinos targeting human beta globulin (5’-CCTCTTACCTCAGTTACAATTTATA-3’) were used as a negative control. Morpholinos were prepared at 200 μM in water, heated for 5 minutes at 65°C and 1 nl (1-2 ng) was injected into the yolk of the zebrafish embryo at the single-cell stage. Injected embryos (morphants) were raised in E3 to 48 hours post fertilization (hpf) and anesthetized with tricaine (168 mg/L) to prepare for live imaging of hydrocephalus (i.e., ventricular enlargement) and otoliths (i.e., number and distance). Images were taken with a Leica MZ75 stereoscope equipped with an Optivision 4K Lite camera. To visualize the developing zebrafish heart and assess laterality defects, mRNA *in situ* hybridization was performed using a protocol modified from (Thisse and Thisse, 2014), with an antisense mRNA probe (414 bp) targeting a cardiac marker *myl7,* synthesized via PCR using cDNA from 24 and 48hpf zebrafish as template, and mRNA synthesis (F: 5’-GACCAACAGCAAAGCAGACA-3’, R: 5’-<u>TAATACGACTCACTATAGGG</u>GGGTCATTAGCAGCCTCTTG-3’, T7 polymerase binding site is underlined). The probe was synthesized using a T7 kit paired with a digoxigenin labeling mix (Invitrogen, AM1334; Roche, 11277073910). At 48 hpf the embryos were fixed in 4% paraformaldehyde for 2h at room temperature. Embryos were washed 3 times for 5 minutes in PBS and stored at −20°C in 100% methanol. On day 1 of the *in situ* hybridization protocol, embryos were rehydrated stepwise in PBS and incubated with the antisense probe at 70°C overnight. On day 2, embryos were washed twice with 20×SSC (3M NaCl, 0.3M trisodium citrate) for 30 minutes at 70°C, and then washed twice for 30 minutes in 2×SSC buffer at room temperature. All solutions were prepared with DEPC-treated water. Embryos were placed in a blocking solution: 5% normal sheep serum, 1% bovine serum albumin fraction V in PBT (PBS with 0.1% Tween 20) for 1 hour at room temperature, and then incubated with 1:1000 anti-digoxigenin-AP (Roche, 11093274910) in the blocking solution for 2 hours. Embryos were then transferred to PBT, and incubated at 4°C overnight. On day 3, embryos were washed 2 times for 15 minutes in staining buffer (100 mM NaCl, 100 mM Tris-HCl). The embryos were then incubated in the NBT/BCIP solution diluted in staining buffer at 1:100 (Roche, 11681451001) and transferred to a 24-well plate for signal development. Once the signal had developed, embryos were post-fixed in 4% PFA for 20 minutes at room temperature and washed 3 times in PBS for 5 minutes. The prepared embryos were then embedded in 3% methylcellulose and imaged from an anterior view.

### Fluorescence Microscopy

For immunofluorescence, vegetatively growing or cilia-regenerating (after a pH shock deciliation (Calzone and Gorovsky, 1982)), *Tetrahymena* cells were fixed using 2% paraformaldehyde and 0.1% Triton X-100 in PHEM buffer, air dried and probed with antibodies as described in (Gaertig et al., 2013). The specificities and dilutions of antibodies used are as follows: anti-polyglycylated tubulin antibodies (polyG, 2302 rabbit polyclonal serum, 1:200 (Duan and Gorovsky, 2002)), anti-polyglycylated tubulin antibody (AXO49, mouse monoclonal, 1:400 (Levilliers et al., 1995)), anti-centrin antibody (20H5, mouse monoclonal, 1:100, EMD Milipore (Manandhar et al., 1999)), anti-acetyl-K40 α-tubulin (6-11B-1, mouse monoclonal, 1:200, Sigma-Aldrich T6793, (Piperno and Fuller, 1985)), anti-GFP antibody (Abcam #6556 or Rockland #600-401-native promoter) were prepared using high-pressure freezing and freeze substitution (Bayless et al., 2016; Dahl and Staehelin, 1989; Meehl et al., 2010). SPEF1B-mCherry was detected by rabbit anti-mCherry antibodies followed by anti-rabbit secondary antibodies conjugated to 15-nm gold particles. Sixty nm sections were imaged on a CM10 electron microscope (Philips) with a BioScan2 CCD camera (Gatan). For immunogold analysis of whole mounts of axonemes, GFP-SPEF1B rescue cells were grown to a density of 2 x 10^5^ cells/ ml, washed with 10 mM Tris-HCl, pH 7.4 and suspended in 20 ml of 10 mM Tris-HCl, 50 mM sucrose, 10 mM CaCl_2_, pH 7.4, deciliated by addition of 420 μl acetic acid, followed after 1 minute by neutralization with 360 μl of KOH. Centrifugation at 1000 x g for 5 min was used to collect the cell bodies. The supernatant was spun at 22000 x g for 20 min at 4 °C to collect cilia. The cilia pellet was suspended in 1.5 ml of motility buffer (1 mM DTT, 50 mM potassium acetate, 5 mM MgSO_4_, 1 mM EGTA, 30 mM HEPES, 1% polyethylene glycol, pH 7.6) and spun again at 22000 x g for 7 min at 4 °C. The pellet was then suspended in 100 μl of motility buffer and 500 μl of 1% NP-40 was added to demembranate. The solution was kept on ice for 15 min. Axonemes were collected by centrifugation at 22000 x g for 7 min at 4 °C, suspended in 100 μl of motility buffer at room temperature and reactivated with 0.5 μl of 100 mM ATP. Ten μl of reactivated axonemes were immediately placed on poly-L-lysine-coated nickel grid (FCF400-Ni, Electron Microscopy Sciences) and allowed to settle for 3 min. The excess liquid was wicked off using a filter paper. The sample was fixed by inverting the grid onto a 10 μl drop of 2% paraformaldehyde and incubating for 15 min. The grids were washed by placing them onto 10 μl drops of PBS (3 x 5 min) followed by an incubation in 10 μl of blocking solution (3% BSA, PBS, 0.01% Tween 20) for 15 minutes. Next, grids were incubated with the anti-GFP antibodies (Abcam6556, rabbit polyclonal, 1:100 in PBS), washed with PBS (5 min) and incubated with secondary antibodies (goat anti-rabbit-10 nm gold, EMS, 1:20) for 1 hour at room temperature. The grids were washed twice with PBS and once with water. The sample was negatively stained with 2% uranyl acetate, air dried and imaged in a JEOL JEM1011 electron microscope.

### Cryo-electron tomography

The dataset for wild-type (WT) *Tetrahymena* cilia with intact membranes was prepared and collected as described previously (Legal et al., 2023). The SPEF1AB-KO cilia were processed using a similar protocol. In brief, cilia were lightly crosslinked with 0.15% glutaraldehyde for 30 minutes on ice, followed by quenching with 35 mM Tris buffer (pH 7.5) and mixing with 5 nm gold beads. A 4 µl aliquot of the crosslinked sample was deposited onto the grid and incubated for 45 sec at 23°C under 100% humidity. The sample was then blotted at force 0 for 8 seconds before being plunge-frozen in liquid ethane. Tilt series were acquired using a 300 kV Titan Krios electron microscope (Thermo Fisher Scientific) equipped with a BioQuantum energy filter (Gatan, Inc.) and a K3 Summit direct electron detector (Gatan, Inc.). Data collection was performed in SerialEM (Mastronarde, 2005) with a grouped dose-symmetric scheme, covering a tilt range of −60° to +60° in 3° increments. Wild-type and SPEF1AB-KO cilia were imaged at 26,000× (pixel size: 3.37 Å) and 42,000× (pixel size 2.12 Å). Each tilt series had a defocus range of −2 to −4 µm and a total cumulative dose of 164 e⁻/Å². Per tilt view, 10-frame movies were recorded. Frame alignment was performed in Warp, followed by tilt-series alignment and tomographic reconstruction in IMOD. Tomograms were denoised by IsoNET for visualization.

### Mass spectrometry

The samples of *Tetrahymena* cilia prepared for cryo-EM were in parallel analyzed by mass spectrometry. Twenty-five to 30 μg of proteins were loaded on an SDS-PAGE gel and subjected to a brief electrophoresis until the samples entered the resolving gel. The gel bands were cut, subjected to reduction with DTT, alkylated with iodoacetic acid and digested with trypsin. Extracted peptides were re-solubilized in 0.1% aqueous formic acid, loaded onto a Acclaim Pepmap (ThermoFisher, 75 μm × 20 mm C18 3 μm beads) precolumn and then separated on an Acclaim Pepmap Easyspray analytical column (ThermoFisher, 75 μm × 15 mm with 2 μm C18 beads) using a Dionex Ultimate 3000 uHPLC at 250 nL/min with a gradient of 2–35% organic solvent (acetonitrile with 0.1% formic acid) over 3 h. Peptides were analyzed using a Orbitrap Fusion mass spectrometer (ThermoFisher) operating at 120,000 resolution (FWHM in MS1) with HCD sequencing (15,000 resolution) at top speed for all peptides with a charge of 2+ or greater. The raw data were converted into the mgf format for searching using the Mascot 2.6.2 search engine (Matrix Science) against the *Tetrahymena thermophila* proteins from UniProt. Mass spectrometry data were analyzed by Scaffold_5.0 (Proteome Software Inc.). Proteins with mean values of exclusive unique peptide count of 2 or more in the wild-type mass spectrometry results were used for analysis. Raw mass spectrometry data were normalized by total spectra. T-test was applied to wild-type and SPEF1AB-KO mass spectrometry results using three biological replicates and p < 0.05 was considered as significant. The SPEF1AB-KO samples were analyzed in the same experiment as already published FAP256AB-KO samples and shared the same wild-type control (CU428) (Legal et al., 2023).

## Movie legends

**Movie 1.** A high-speed DIC video of wild-type cell swimming inside a microfluidic channel.

**Movie 2**. A high-speed DIC video of SPEF1AB-KO cell swimming inside a microfluidic channel. Green arrow points to a cilium that shows an abnormal shape of the recovery stroke. Yellow arrows point to cilia that occasionally stall. Red arrow shows a cilium with an incomplete power stroke.

**Movie 3. TIRFM movie showing the localization of mNeonGreen-SPEF1B on cortical microtubules.** Yellow arrows; longitudinal microtubules, magenta arrows; post-ciliary microtubules, red arrows; transverse microtubules, blue arrow; contractile vacuole pore microtubules.

**Movie 4. SPEF1 localizes to the cortical microtubules.** A two-color TIRFM movie reveals SPEF1B along the longitudinal and transverse microtubule bundles (yellow arrows) in cells expressing mNeonGreen-SPEF1B and β-tubulin-mCherry (red).

**Movie 5. SPEF1 is concentrated at the distal ends of cilia.** A two-color TIRFM movie reveals SPEF1B concentrated at the tips of cilia in the mNeonGreen-SPEF1B (green) and β-tubulin-mCherry (red) expressing cells.

**Movie 6. Imaging of mNeonGreen-SPEF1B in mature cilia.** A TIRFM video of a SPEF1AB-KO cell rescued by a transgene expressing mNeonGreen-SPEF1B. See the kymographs of cilia in Figure 6A.

**Movies 7. SPEF1 turns over slowly at the distal ends of mature cilia.** A FRAP assay was performed on the cilium marked with an arrow in the cell expressing mNeonGreen-SPEF1B. The signal quantifications in the bleached (arrow) and control unbleached cilium on the left are shown in Figure 6B.

**Movie 8. SPEF1 forms stationary particles at the tips of assembling cilia.** A TIRFM movie of mNeonGreen-SPEF1B at the tips of assembling cilia. The cell was deciliated with a pH shock, and imaged 2 hr later as cilia regrow. Multiple tips of assembling cilia are pressed against the glass as the cell is trapped inside a small volume. The kymographs of several cilia documenting the positions of SPEF1B particles are shown in Figure 6C.

**Movie 9. SPEF1B turnover slowly at the tips of assembling cilia.** A TIRFM movie of mNeonGreen-SPEF1B at the tips of assembling cilia. The cell was deciliated with a pH shock, and imaged 2 hr later as cilia regrow. A FRAP assay was performed on the cilium marked with an arrow in the cell expressing mNeonGreen-SPEF1B. The signal quantifications in the bleached (arrow) and control unbleached cilium on the left are shown in Figure 6D.

## Supplementary figures

**Figure S1.**
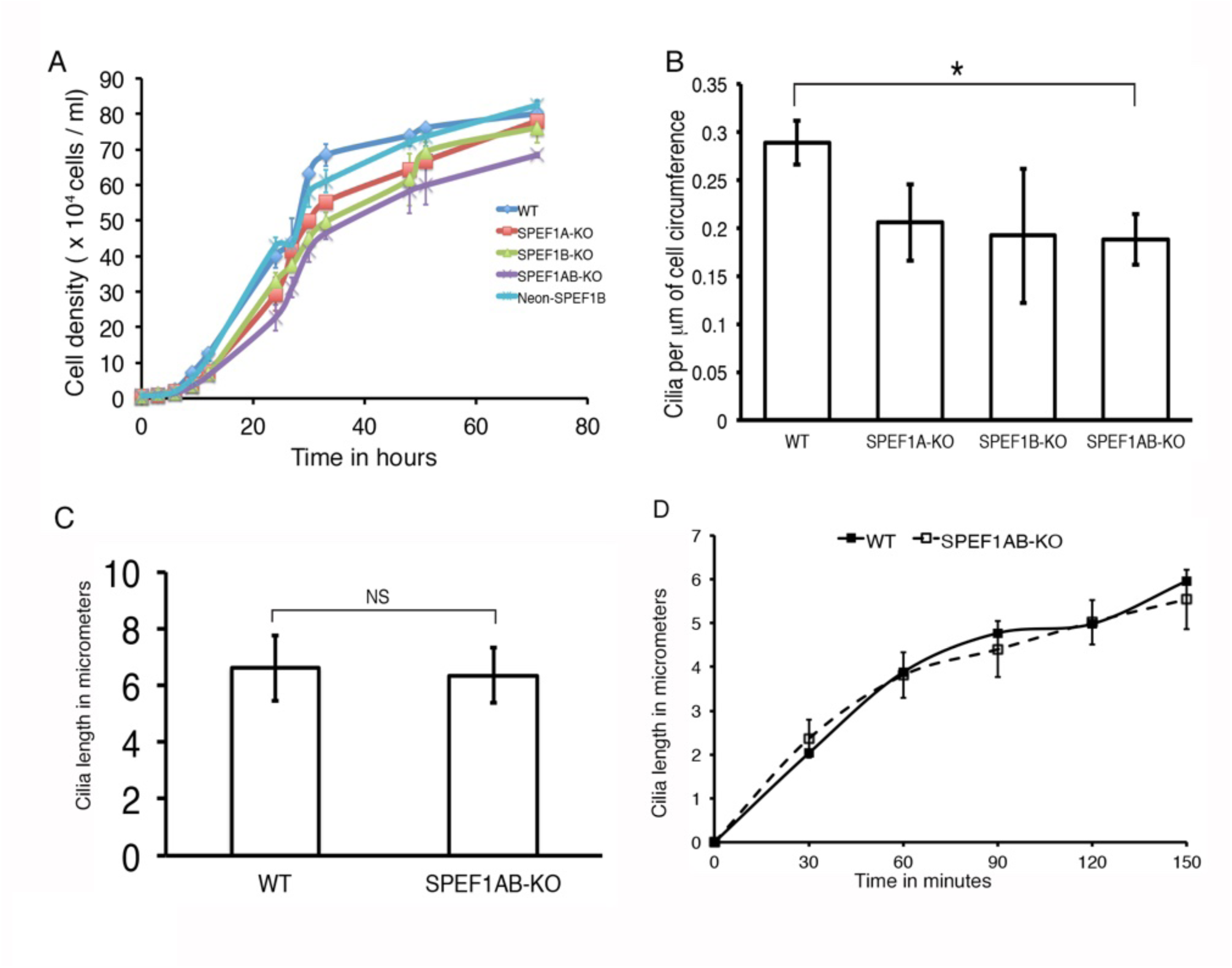
Quantifications of cilia length, rate of cilia regeneration, cilia density and rate of cell multiplication. (A) Culture growth rates based on mean cell densities from two independent experiments. (B) Cilia density quantified as the number of cilia per mm of cell circumference (at the widest cell circumference). Mean ± SD. WT: N = 5 cells; SPEF1A-KO: N = 5 cells; SPEF1B-KO: N = 5 cells and SPEF1AB-KO: N = 14 cells. The scores represent the number of cilia visible at the widest cell circumference in a confocal section by the length of the circumference. *P< 0.0001 (two tailed t test). (C) Average cilium length. Mean ± SD μm. WT: 6.62 ± 1.16, N = 107 cilia, N = 6 cells. SPEF1AB-KO: 6.36 ± 0.99, N = 104 cilia, N =13 cells. The difference between the length of cilia in wild-type and SPEF1AB-KO cells was not significant (P = 0.07, two tailed t test). (D) The average length of cilia (μm) in cilia-regenerating cells (following deciliation) at different time points. The data were obtained using 5-14 cells per genotype per time-point. Error bars represent standard deviations.

**Figure S2.**
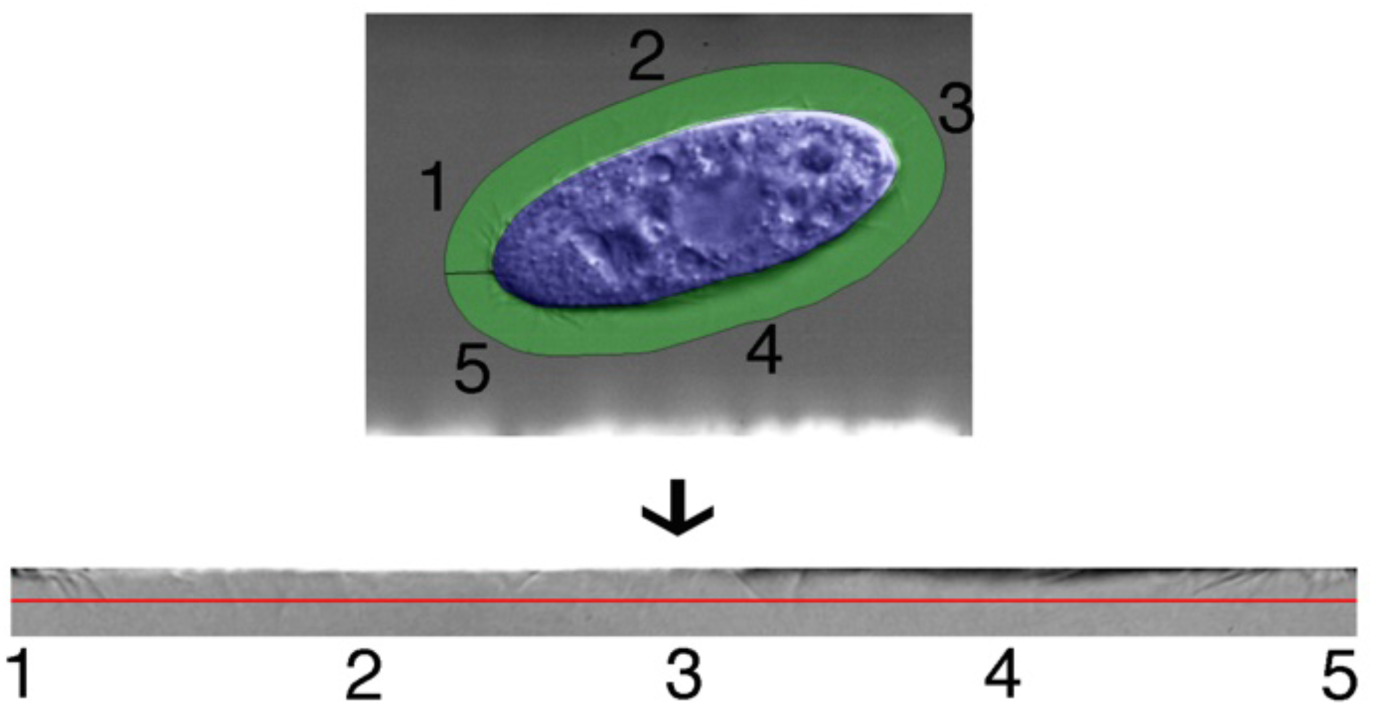
Outline of video processing steps used to generate chronographs (Figure 1F,G) Based on (Funfak et al., 2015; Stoddard et al., 2018). (Top panel) The first frame of a wild type video showing how the cell body (blue) and cilia (green) boundaries were defined. Numbers 1-5 corresponds to the anterior (1 and 5), sides (2 and 4) and posterior (3) areas around the circumference of the cell. (Bottom panel) The first frame of the extracted cilia video that corresponds to the cilia zone marked in the top panel. Numbers 1-5 marks the position of cilia along the circumference of the cell as shown in the top panel. The greyscale was plotted over time at the position marked by the red line to produce a chronograph (this chronograph is shown in Fig. 1F).

**Figure S3.**
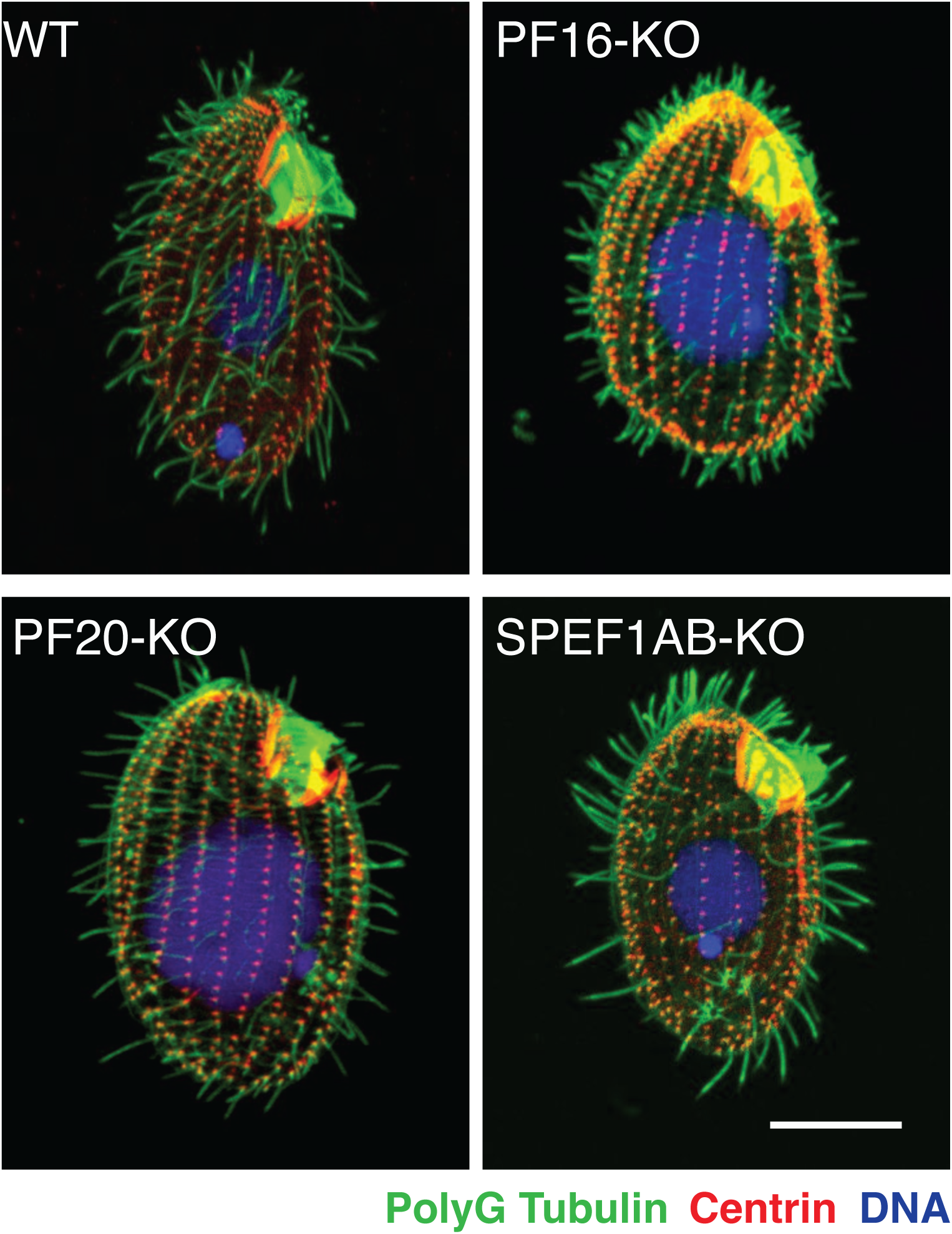
*Tetrahymena* CP mutants have fewer or shorter cilia. Loss of function mutations in the CP proteins, PF16 and PF20, reduce the number of cilia ( as observed in the SPEF1AB-KO cells). In Confocal immunofluoroscence images of WT, PF16-KO, PF20-KO and SPEF1AB-KO cells stained with antibodies against polyglycylated tubulin (green) and centrin (red). Nuclei were stained with DAPI (blue). Scale bar = 10 μm.

**Figure S4.**
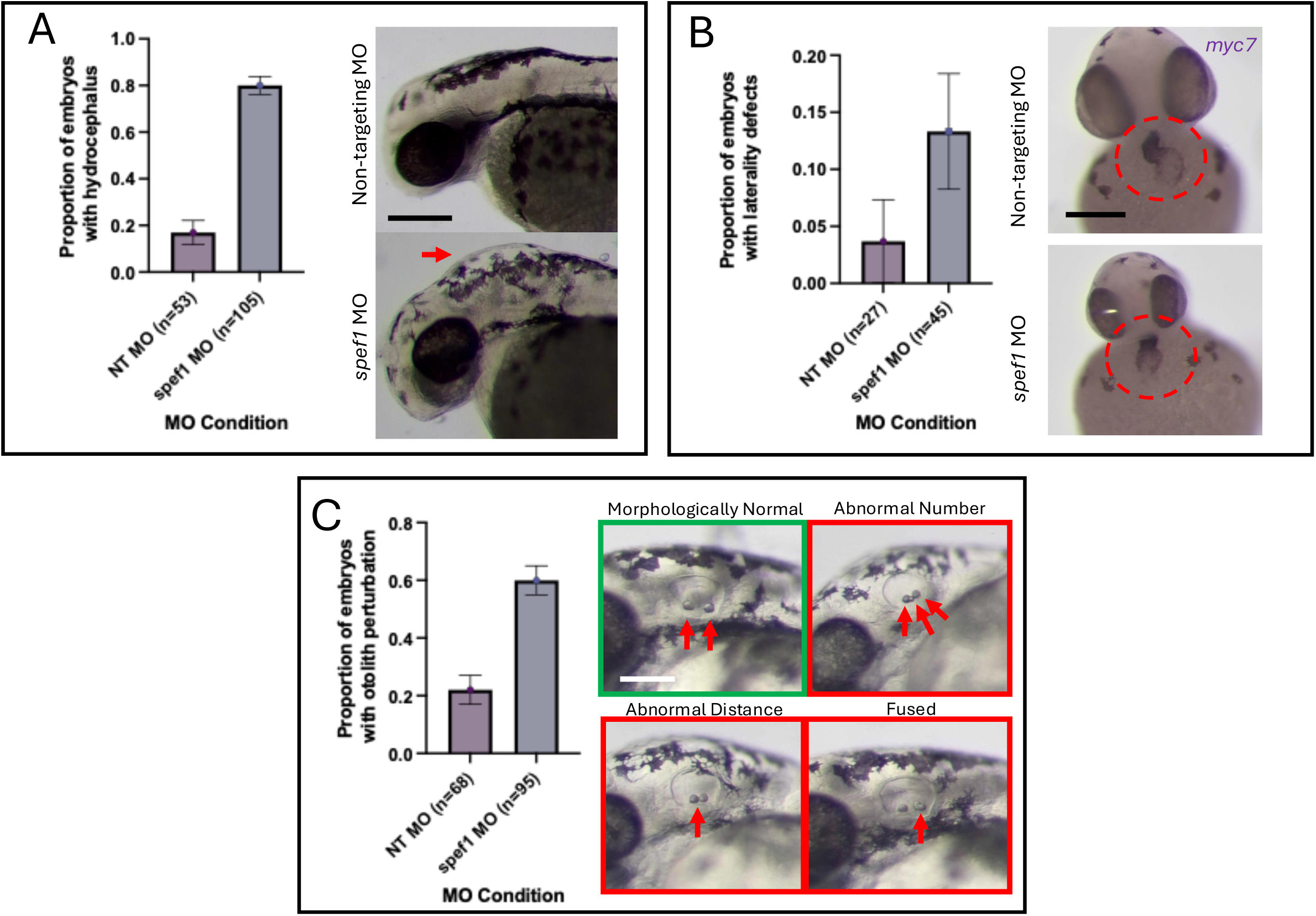
Morpholino-mediated knockdown of *spef1* in 48 hpf zebrafish embryos results in phenotypes consistent with motile cilia dysfunction. (A) *spef1* morphants exhibited a markedly increased incidence of hydrocephalus compared with non-targeting controls (red arrow). Proportions = 16.98% vs. 80.00%. N = 2 biological replicates. Fisher’s exact test: p < 0.0001. 95% binomial CI (morphants) = 0.045–0.176. (B) *spef1* morphants showed an elevated frequency of cardiac laterality defects, assessed by *myl7* in situ hybridization. Proportions = 3.7% vs. 13.33%. N = 1 biological replicate. Fisher’s exact test: p = 0.2440. 95% binomial CI (morphants) = 0.004–0.579. (C) *spef1* morphants displayed abnormal otolith development. Three classes of abnormalities were scored (examples in red boxes): altered otolith number, increased otolith separation, and fused otoliths. Morphologically normal otoliths are shown for comparison (green box). Proportions = 22.06% vs. 60.00%. N = 1 biological replicate. Fisher’s exact test: p < 0.0001. 95% binomial CI (morphants) = 0.121–0.320. Statistical analysis: All statistical comparisons were performed using Fisher’s exact test. Exact 95% binomial confidence intervals (Clopper–Pearson) are reported for morphant proportions in the figure legend. Error bars in the figure represent the standard error of the proportion (SE) and do not correspond to the Clopper–Pearson confidence intervals. Scale bars: ∼100 μm.

**Figure S5.**
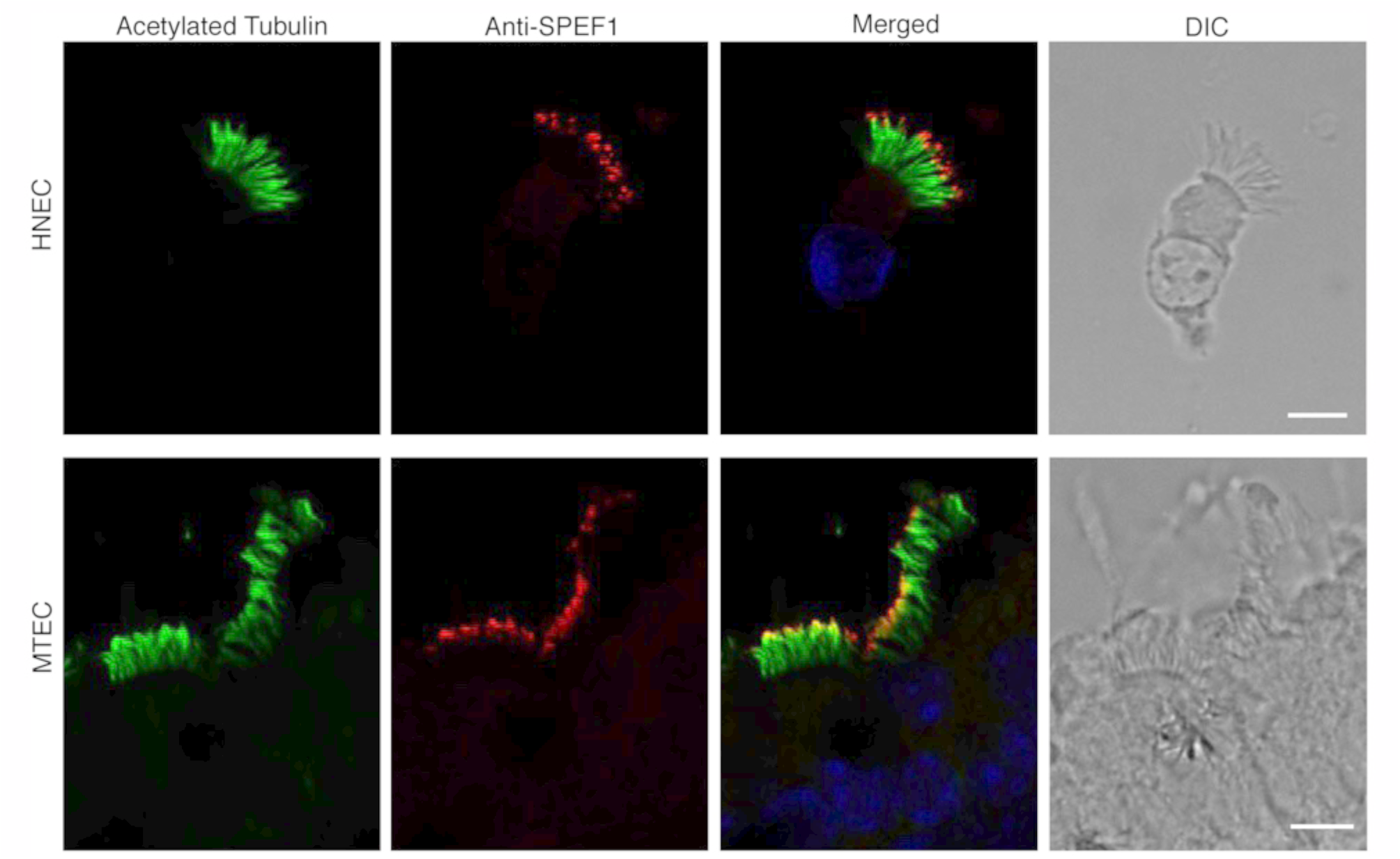
SPEF1 is concentrated near the tips of cilia in mammalian multiciliated cells. Confocal immunofluorescence images of human nasal epithelial cells (HNEC) and murine tracheal epithelial cells (MTEC) obtained using antibodies against acetylated tubulin that highlight axonemes (red) and antibodies against SPEF1 (green). SPEF1 is enriched near the distal tips of cilia. Scale bar = 5 μm.

**Figure S6.**
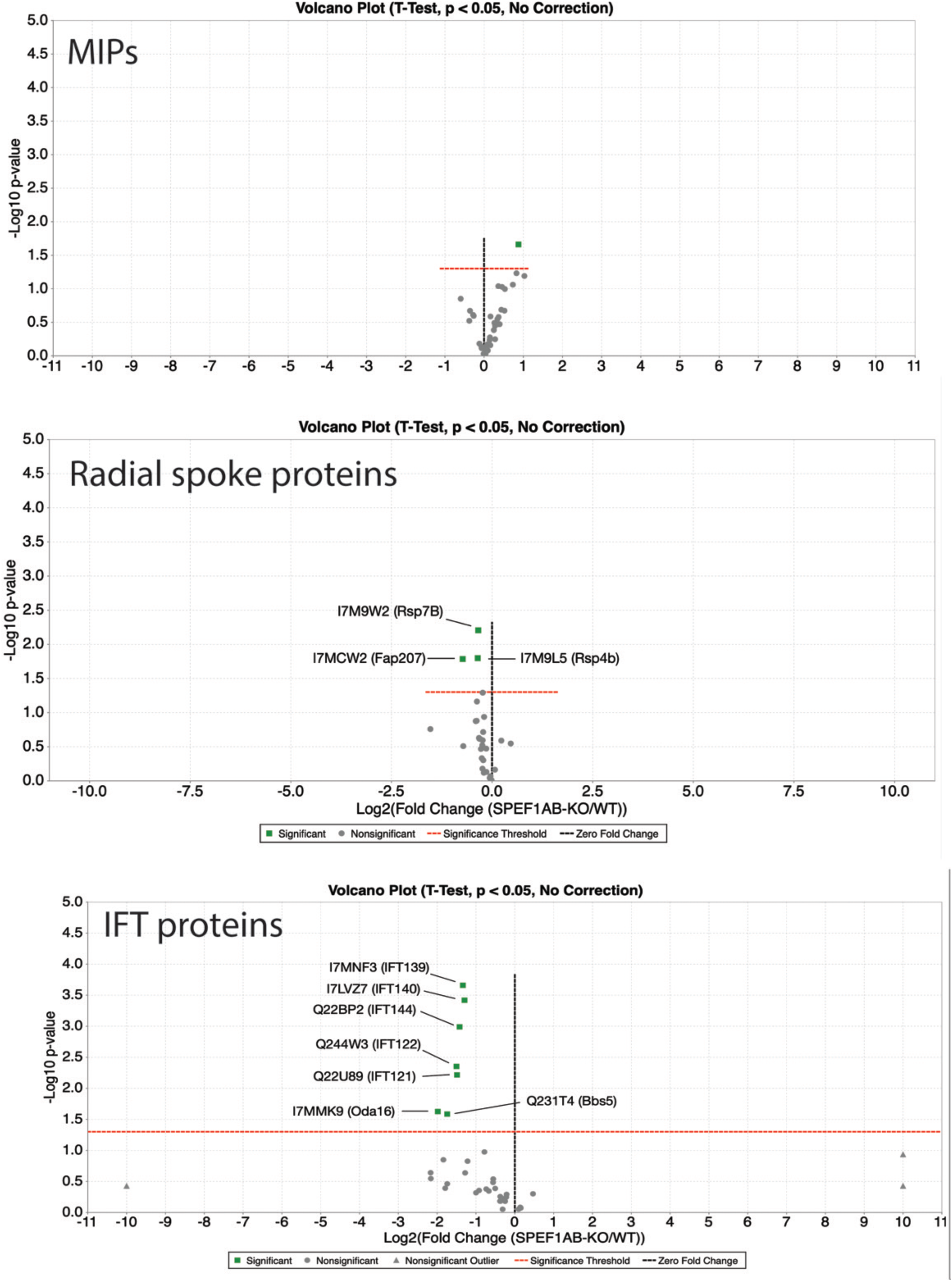
SPEF1 loss does not affect the levels of MIPs and RSPs but depleted IFT complex A proteins. Volcano plots present mass spectrometry-based data comparing the SPEF1AB-KO and wild-type cilia.

**Figure S7.**
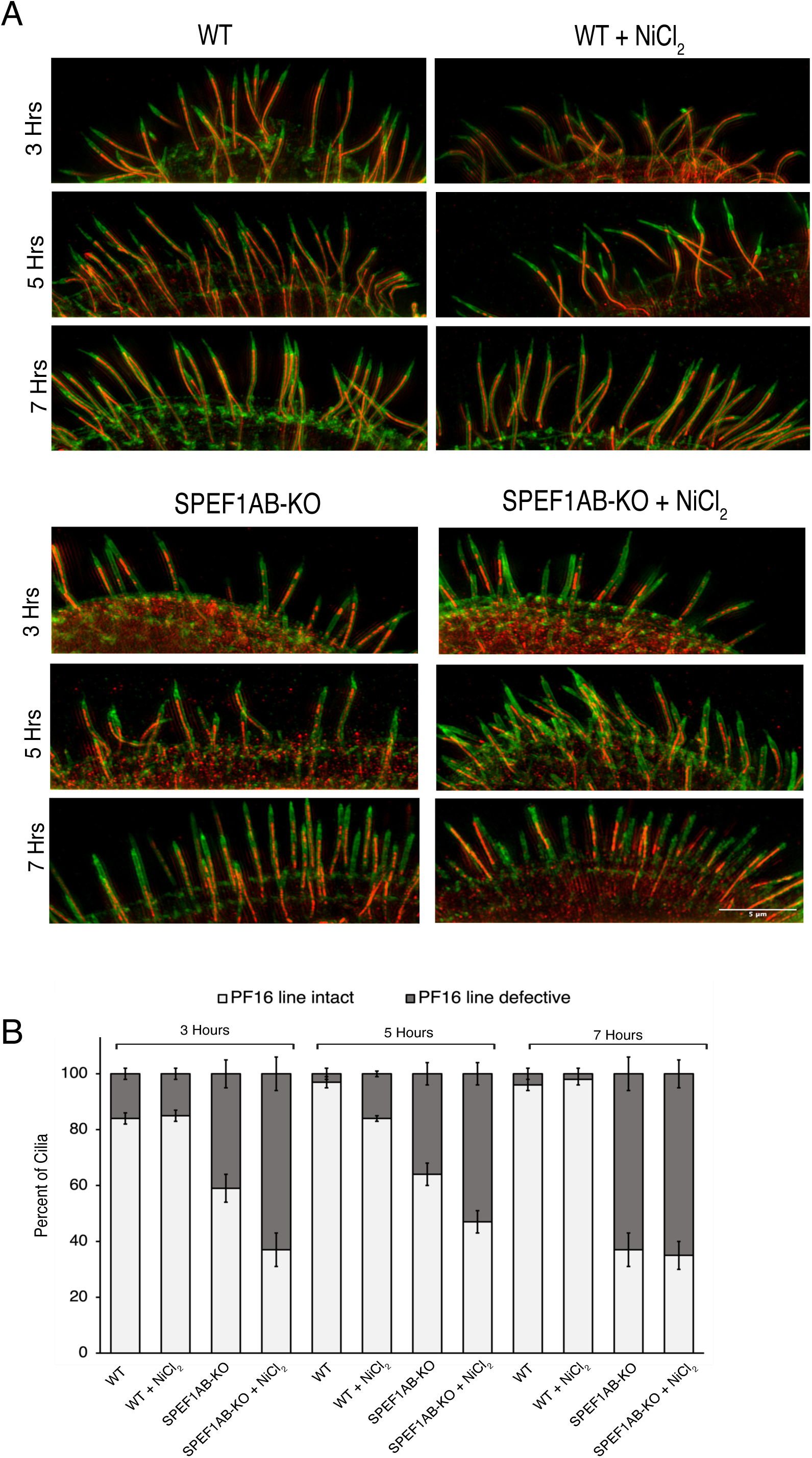
Inhibition of ciliary motility with NiCl_2_ does not rescue the CP defects caused by the loss of SPEF1. (A) SR-SIM images of wild-type and SPEF1AB-KO cells expressing PF16-GFP, which regenerate cilia after deciliation, either in the absence of presence of 500 mM NiCl_2_. Cells were labeled by antibodies agaisnt polyglycylated tubulin (green) and anti-GFP antibodies to detect either PF16-GFP (red). Scale bar = 5 μm. (B) Quantification of the frequency of cilia with a defective PF16-GFP line.

## Supplementary Tables

**Table S1.** Description and sequence of oligonucleotide primers used in this study.

**Table S2.** Analysis of the number of microtubule profiles in cortical bundles in SPEF1AB-KO and wild-type cells.

**Table S3.** Mass spectrometry data for all proteins identified in the wild-type and SPEF1AB-KO cilia (total unique peptide counts and normalized emPAI scores).

**Table S4.** A subset of mass spectrometry data focused on the CP proteins identified in the wild-type and SPEF1AB-KO cilia (total unique peptide counts and normalized emPAI scores).

**Table S5.** A subset of mass spectrometry data focused on the MIPs identified in the wild-type and SPEF1AB-KO cilia (total unique peptide counts and normalized emPAI scores).

**Table S6.** A subset of mass spectrometry data focused on the RSPs identified in the wild-type and SPEF1AB-KO cilia (total unique peptide counts and normalized emPAI scores).

**Table S7.** A subset of mass spectrometry data focused on the IFT proteins identified in the wild-type and SPEF1AB-KO cilia (total unique peptide counts and normalized emPAI scores).

## Acknowledgements

We thank David Mitchell (SUNY Upstate Medical University, Syracuse, NY) for valuable comments and Mary Ard (Georgia Electron Microscopy facility at UGA) for assistance with transmission electron microscopy. The super-resolution imaging was performed at the UGA Biomedical Microscopy Core. Research reported in this publication was supported by the National Institutes of Health including grants: R01GM089912 and R01GM135444 (to J.G.), R01GM110413 and R35GM152057 (to K.F.L.), and R01GM099820 (to C.G.P.). G.W.D. and H.O. were supported by the European Union FP7 projects SYSCILIA (grant 241955) and BESTCILIA (grant 305404) and by the German Research Foundation grants DFG OM6/7 und OL450/1. R.T. and C.N.B. were funded by European Research Council (ERC) grant 278248 Multicell. E.J. was funded by the National Science Centre, Poland, grant no. 024/55/B/NZ3/01651. T.M.K. was funded by the Grand Défi Pierre Lavoie Foundation through the Montreal Children’s Hospital Foundation and the SickKids New Investigator Research Grant (NI21-1159). Z.W.N. was funded by Fonds de Recherche du Québec - Santé (FRQS), Doctoral (347103). K.H.B. was supported by grants from the Canadian Institutes of Health Research (PJT-156354) and the Natural Sciences and Engineering Research Council of Canada (RGPIN-2022-04774). A.G. was supported by the Fonds de Recherche du Québec Nature et Technologies (FRQNT) studentship (345547).

