## Supplementary material for "SPEF1 mediates assembly of the central pair microtubule complexes in cilia of *Tetrahymena*": Table S1

| Description | Restriction sites (Primer 1, Primer 2) | Primer 1 (5’ to 3’) | Primer 2 (5’ to 3’) |
| --- | --- | --- | --- |
| Amplify 5’UTR of SPEF1A | ApaI , SmaI | ATAAGGGCCCAGGTCTTGACGGAGAAGTT | AATTCCCGGGAACCTATTTAGATTCGCTCAA |
| Amplify 3’UTR of SPEF1A | PstI, SacII | AATTCTGCAGTAGATACTTGAACTAAAGATCA | TTATCCGCGGTAAATATATAAATCGTGGTATTC |
| Check deletion of SPEF1A |  | ACTCCTTGTTTAATTTCATATC | AACTTGGGACAATAATGGCT |
| Amplify 5’UTR of SPEF1B | ApaI, Sma | ATTTGGGCCCTCAGATTCAATTTAACATTCAC | AATTCCCGGGTTATAGTTTGTTTCTTTTATCTC |
| Amplify 3’UTR of SPEF1B | PstI, SacII | ATAACTGCAGAGTAAGATGAAATCATTTGGG | TTATCCGCGGTAGCCTTATAGAAAATCATGTT |
| Check deletion of SPEF1B |  | TACAACAAATTATGGAATAAGAA | CTACTACTTCTGCCATTAGA |
| Amplify coding region of SPEF1B (to create GFP-SPEF1B) | MluI, BamHI | AATTACGCGTCATGGAAGAAGCTCCTCATTTA | AATTGGATCCTCAAATCTATGTTAGCTAATTTAT |
| Amplify 5’UTR of PF20 | SacI, BamHI | TTATAGAGCTCGCAACGGGTTACAAGACT | ATATTGGATCCTGGCTTTTTATCTTCCTTAG |
| Amplify 3’UTR of PF20 | XhoI, BamHI | AATTACTCGAGATATCATTTATCCTTGCTTCTA | TAATTGGATCCCGAAGATAAAGTAGAAGACG |

Supplementary Table 1
